# Sheathless elasto-inertial focusing of sub-25 nm particles in straight microchannels

**DOI:** 10.1101/2025.02.16.638527

**Authors:** Selim Tanriverdi, Javier Cruz, Shahriar Habibi, Taras Sych, Martim Costa, Gustaf Mårtensson, André Görgens, Samir EL Andaloussi, Luca Brandt, Outi Tammisola, Erdinc Sezgin, Aman Russom

## Abstract

Nanoscale biological particles, such as lipoproteins (10–80 nm) or extracellular vesicles (30–200 nm), play pivotal roles in health and disease, including conditions like cardiovascular disorders and cancer. Their effective analysis is crucial for applications in diagnostics, quality control, and nanomedicine development. While elasto-inertial focusing offers a powerful method to manipulate particles without external fields, achieving consistent focusing of nanoparticles (<500 nm) has remained a challenge. In this study, we experimentally demonstrate elasto-inertial focusing of nanoparticles as small as 25 nm using straight high-aspect-ratio microchannels in a sheathless flow. Systematic investigations reveal the influence of channel width, particle size, viscoelastic concentration, and flow rate on focusing behavior. Additionally, through numerical simulations and experimental validation, we provide insights into particle migration dynamics and viscoelastic forces governing nanoparticle focusing. Finally, we successfully focus biological particles, including liposomes (90–140 nm), extracellular vesicles (100 nm), and lipoproteins (10–25 nm), under optimized conditions, showcasing potential applications in medical diagnostics and targeted drug delivery. These findings mark a significant advancement toward size-based high-resolution particle separation, with implications for biomedicine and environmental sciences.

## Background

The ability to manipulate nanoparticles is crucial for many fields ranging from disease diagnostics to drug delivery.^1^ Nanoscale bioparticles in the body play an indispensable role in health and their dysregulation causes many diseases. For example, lipoproteins, such as high-density lipoprotein (HDL) and low-density lipoprotein (LDL), transfer lipids through the body. Their imbalance, as well as their deviation from canonical physical properties such as size, is a marker of dyslipidemia and metabolic diseases.^2^ Moreover, certain types of lipoproteins are associated with neurodegenerative diseases such as Alzheimer’s.^3,4^ Extracellular Vesicles (EVs) enable intercellular communication in the body, and certain types of EVs are associated with diseases like cancer.^5^ Current methods to sort biological nanoparticles, such as size exclusion chromatography (SEC),^6^ asymmetrical flow field-fractionation (AF4)^7^ or centrifugal techniques^8^ are cumbersome, expensive, and rarely result in pure fractions. Therefore, developing advanced methods to sort and study particles based on their physical properties, such as size, is crucial.

In addition to biomedical applications involving biological nanoparticles, environmental nanoparticles are also relevant for human health. For example, microplastics in water pose a significant threat to human health. The toxicity and environmental impact of these contaminants is highly affected by particle size and properties.^9^ Therefore, size-based enrichment of nanoparticles is critical for downstream analysis in environmental studies towards determining the toxicities of these particles in solution.^10^

Microfluidics has emerged as a promising technology over the last few decades, revolutionizing fields ranging from biomedicine to chemistry and environmental science.^11–13^ The ability to manipulate fluids at the microscale provides unparalleled control over particle sorting, focusing, and separation, making microfluidics a powerful tool for research and industrial applications.^14^ Microfluidic manipulation methods are broadly categorized into active and passive methods, depending on whether external forces are used to manipulate particles. Active methods rely on external forces, such as electric,^15^ magnetic,^16^ or acoustic^17,18^ waves to direct particle movement. These methods offer high precision micrometer sized particles but often require complex setups and significant operational costs. In contrast, passive methods utilize inherent fluid properties and microchannel geometry to achieve particle manipulation without external actuation, making the methods simpler and more cost-effective. Among passive methods, inertial microfluidics^19^ and elasto-inertial microfluidics^20^ have garnered significant attention due to their ability to perform label-free focusing and separation at high-throughput.

Inertial microfluidics leverages the interplay of shear-induced lift forces and wall-induced lift forces to focus particles at equilibrium positions within Newtonian fluids.^21,22^ A single, stable focusing position^23,24^ and high resolution separation^25^ can be achieved by the addition of curvature, but the systems demand very high pressures (tens to hundreds of bar) to focus sub-micron particles, leaving nanoparticles out of reach. Elasto-inertial microfluidics extends the principles of inertial microfluidics by introducing viscoelastic fluids, which generate an additional elastic force arising from the fluid’s normal stress differences.^26^ The combination of elastic, shear-induced, and wall-induced lift forces enables the focusing of particles at a single, stable equilibrium position,^27^ even for submicron particles. This enhanced control makes elasto-inertial microfluidics particularly suitable for nanoparticle applications. One important advantage of elasto-inertial focusing is its enhanced control over submicron particles, eliminating the effect of Brownian motion, which becomes more significant as particle size decreases.^28^

Despite the promise of elasto-inertial microfluidics, its application to nanoparticle manipulation remains underexplored. Previous studies have achieved focusing of particles down to 100 nm in curved, spiral microchannels^29^ and 200 nm in straight microchannels.^30^ However, focusing particles smaller than 100 nm in straight microchannels has remained a challenge due to the high pressures required for effective manipulation.

In this study, we experimentally investigate elasto-inertial focusing of nanoparticles as small as 25 nm using high-aspect-ratio microchannels in sheathless flow conditions. Numerical simulations complement these experiments by providing detailed insights into the forces and dynamics governing particle migration towards the equilibrium position. We systematically evaluate the effects of particle size, channel geometry, viscoelastic fluid properties, and flow rates on focusing behavior. Our experiments include the use of polystyrene beads (25, 50, 100, 200, 500 nm and 1 µm), two different channel widths (5 and 10 µm) and four viscoelastic fluid concentrations (500, 1000, 2000 and 4000 ppm of PEO). Additionally, we demonstrate the successful focusing of biological nanoparticles, including liposomes (90 – 140 nm), extracellular vesicles (100 nm), high and low–density lipoproteins (10-25 nm), at the optimal experimental conditions. These findings lay the foundation for advanced nanoparticle manipulation strategies and their applications in biomedicine and environmental sciences.

## THEORETICAL BACKGROUND

Elasto-inertial microfluidics relies on a detailed understanding of the flow regime and the forces acting on particles within a microchannel. To better understand how these forces affect particle behavior under different flow conditions, the use of dimensionless numbers such as the Reynolds number (Re), Weissenberg number (Wi) and Elasticity number (El) are useful. Re describes the ratio of inertial to viscous forces in a fluid and determines the flow regime and is formulated as^21^ *Re = ρU_m_L/µ*, where *ρ* is fluid density, *U_m_* is the average fluid velocity, *L* is the characteristic channel length, and *µ* is the fluid viscosity. The Weissenberg number quantifies the ratio of elastic to viscous effects in a viscoelastic fluid, providing insight into the role of fluid elasticity on particle behavior. Wi is expressed as^31^ *Wi = 2λQ/hw^2^*, where *λ* is the relaxation time of the viscoelastic fluid, *Q* is the volumetric flow rate, *h* is the height of the channel and *w* is the channel width. The importance of the Weissenberg number arises as it involves the characteristic relaxation time of the polymers, which is an intrinsic property of non-Newtonian fluids and varies with the polymer concentration of the fluid. This parameter is crucial for understanding elastic contribution to particle migration. Finally, the Elasticity number combines Re and Wi to describe the relative importance of elastic and inertial effects. It is defined as (*El = Wi/Re*)^31^ and is instrumental in interpreting experimental data, as it reflects the balance between inertial and elastic forces.

The blockage ratio is another dimensionless number that plays a role in particle focusing. It is defined as the ratio of particle size over characteristic dimension of the channel (*ß = a/L*).^32^ In this study, we define the blockage ratio as *ß = a/w*, since the microchannel width represents the characteristic length of the high aspect ratio microchannels (*AR=h/w*) considered in this study.

Elasto-inertial focusing occurs when the lift force and the elastic force balance each other in a microchannel. The lift force (*F_L_*) is considered as the combination of shear-induced lift force and wall-induced lift force and formulated as^33^ *F_L_ = c_L_ρU ^2^a^4^/D_h_ ^2^*, where *c_L_* is the lift coefficient, *a* is the particle size, and *D_h_* is the hydraulic diameter of the channel. The presence of non-Newtonian fluid in a microchannel causes unequal normal stress differences (*N_1_* and *N_2_*), which results in the formation of an elastic force.^34^ The elastic force is expressed as^35^ *F_E_ = a^3^∇ N_1_*. The final equilibrium position of the particles depends on the complex relation between the lift force and the elastic force, and the geometry of the cross-section of the microchannel. In previous studies, it has been shown that particles can be found at the channel corners or at the center in single or multiple positions depending on the flow conditions and channel geometry.^20^

## RESULTS AND DISCUSSION

In this section, we present experimental and numerical results on nanoparticle focusing and migration in elasto-inertial microfluidics. First, we compare inertial and elasto-inertial focusing for 200 nm, 500 nm, and 1 µm particles. Then, we systematically investigate the effects of particle size, channel width, viscoelastic fluid concentration, and flow rate on focusing behavior. Additionally, we explore particle migration dynamics using numerical simulations to understand the interplay between elastic and inertial forces during size-based migration toward equilibrium positions. Finally, we demonstrate the potential of this method for biological nanoparticle focusing under optimized conditions, paving the way for applications in diagnostics, particle enrichment. And high-resolution size-based separation.

### Inertial Focusing vs. Elasto-inertial Focusing

The ability to focus nanoparticles is a critical challenge in microfluidics, primarily because the forces governing particle migration scale with particle size. As it will be described below, this is the case particularly when comparing inertial and elasto-inertial approaches. As shown schematically in Figure 1(a), inertial microfluidics typically focuses particles along the channel’s center-face, resulting in two equilibrium positions in high-aspect-ratio straight channels. However, when viscoelastic fluids, such as PEO solutions, are used in elasto-inertial microfluidics, the addition of elastic forces drives particles to focus at a single central position, fundamentally altering their behavior. Experimental results for 200 nm, 500 nm, and 1 µm particles under identical flow rates (3 µL/min) and channel geometry (*h = 60 µm, w = 5 µm*) are shown in Figure 1(b). In Newtonian fluids, 1 µm particles exhibit partial focusing near the two long sides of the microchannel, as expected in inertial focusing within high-aspect-ratio channels.^21,36^ However, this focusing is incomplete, and smaller particles (200 nm and 500 nm) show no focusing behavior due to insufficient inertial forces,^37^ emphasizing the size dependency of inertial focusing. Achieving nanoparticle focusing with inertial microfluidics would require impractically high flow rates, making this approach unsuitable for such applications.

**Figure 1.**
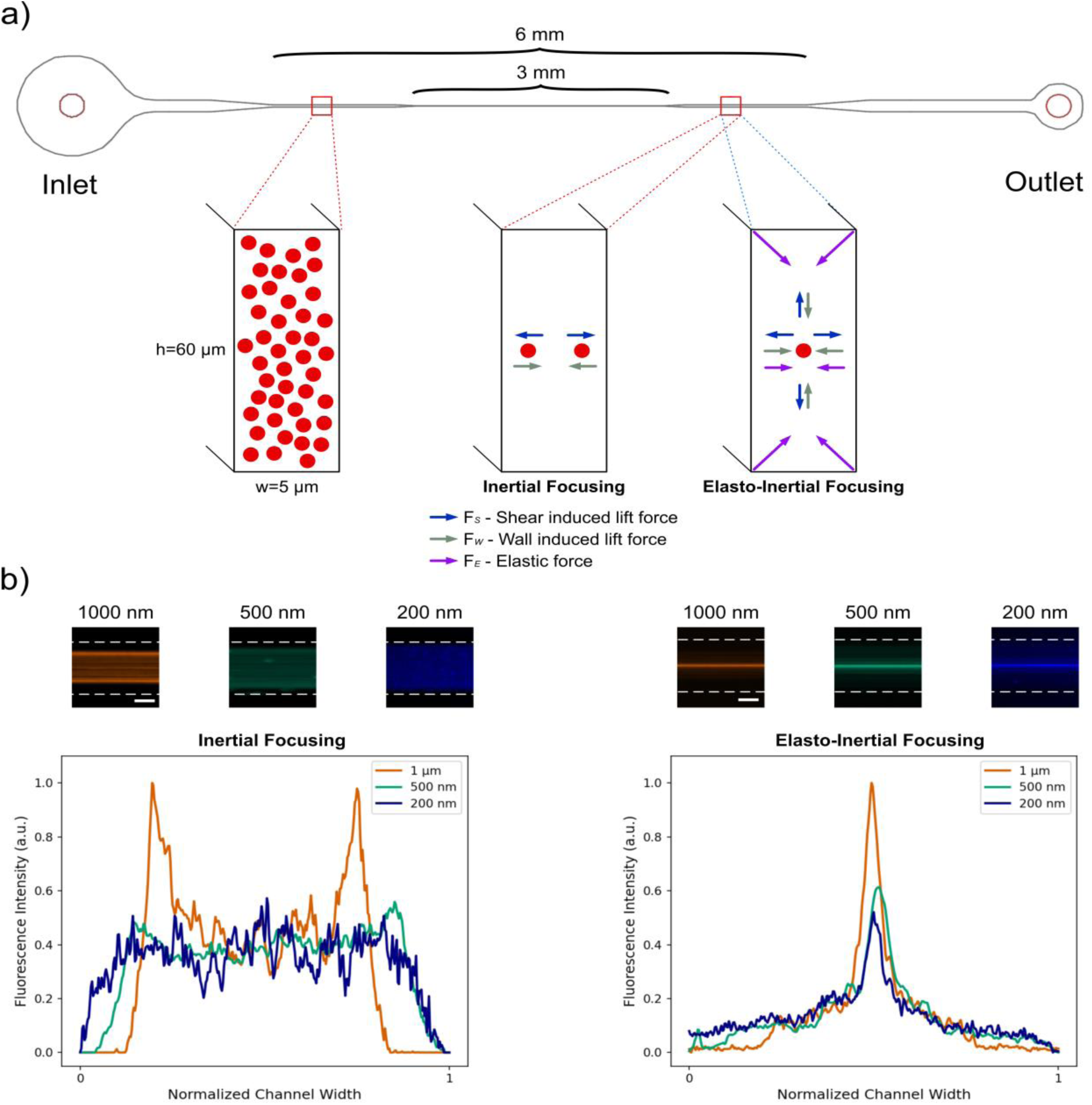
Overview of particle focusing principle in high-aspect-ratio microchannels. a) Schematic illustration of the microfluidic device, showing the design and the cross-sectional view of the microchannel. Inset highlights the high-aspect-ratio segment and forces involved in inertial and elasto-inertial focusing. b) Inertial focusing (left) and elasto-inertial focusing (right) in a high-aspect-ratio microchannel (*h=60 µm, w=5 µm*). The upper panel shows the fluorescence images of 1 µm (orange), 500 nm (green), and 200 nm (blue) particles, while the lower panel presents the corresponding cross-sectional intensity profiles. The flow rate is 3 µL/min. Scale bar: 50 µm.

In contrast, elasto-inertial microfluidics using a 1000 ppm PEO solution demonstrates successful focusing for all three particle sizes at the channel center, with 1 µm particles displaying the highest fluorescence intensity. These results highlight the size-dependent nature of elasto-inertial focusing,^38^ where elastic forces significantly enhance the manipulation of smaller particles. Importantly, this study demonstrates that elasto-inertial microfluidics overcomes the limitations of inertial focusing, enabling nanoparticle alignment under practical experimental conditions.

### Achieving Particle Focusing Down to 25 nm

The ability to focus sub-100 nm particles represents a significant milestone in elasto-inertial microfluidics, addressing long-standing challenges in manipulating nanoscale particles effectively. After successfully focusing 200 nm and 500 nm particles, we extended our investigation to particles ten times smaller. Figure 2 shows the focusing behavior of 25 nm particles under a fixed PEO concentration of 500 ppm in a microchannel with an aspect ratio of 6 (*h=60 µm, w=10 µm*), at four different flow rates (0.5, 1, 1.5 and 2 µL/min).

**Figure 2.**
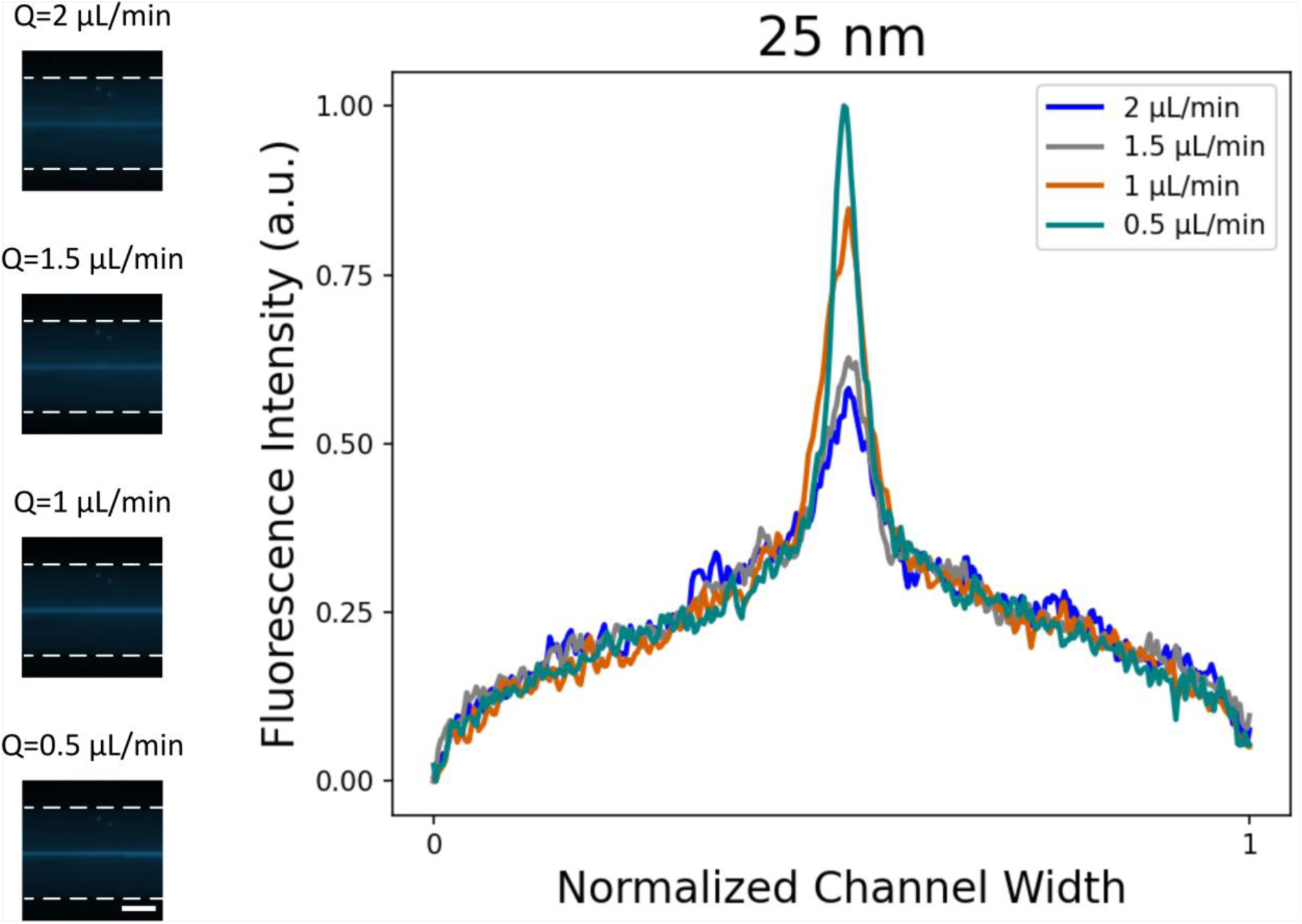
The focusing of 25 nm particles. The graph shows the normalized intensity of the cross-section of particle focusing in a high-aspect-ratio microchannel (*h=60 µm, w=10 µm*) at flow rates ranging from 0.5 to 2 µL/min. The inset shows the corresponding fluorescence microscopy images of 25 nm nanoparticles focusing at varying flow rates. Scale bar: 50 µm.

These experimental conditions (500 ppm of PEO and channel w=10 µm) were chosen to identify the lower limit of polymer concentration required to maintain focusing behavior at larger channel widths. Fluorescence intensity graphs reveal that 25 nm particles predominantly focus at the channel center, albeit with slight distortions. The highest focusing quality was observed at the lowest flow rate (0.5 µL/min), with quality gradually decreasing as flow rates increased due to reduced residence time hindering particles migration fully to the center. With a blockage ratio (a/w) of 0.0025, these results are approximately 20 times smaller than previously reported focusing thresholds, demonstrating the dominant role of viscoelasticity (Re<1, Wi>10, El≈63) in overcoming Brownian motion and achieving precise nanoparticle focusing.

To achieve such high focusing efficiency without significant pressure drops, we implemented an innovative microchannel design previously introduced by our team.^39^ This design features a short segment with a small channel width, creating a high-aspect-ratio that enhances focusing. Following this segment, the channel widens, reducing overall resistance and preventing significant pressures increases within the system. This configuration ensures that particles maintain their focused positions at the channel center even after transitioning into the wider section.

By implementing this design, we effectively balance the benefits of high-aspect-ratio focusing with manageable pressure drops, enhancing the practicality of elasto-inertial microfluidic systems. We achieve precise nanoparticle focusing without the drawbacks associated with uniformly narrow channels, such as excessive pressure drops. Building on these results, we next investigate the influence of channel geometry and fluid properties on the focusing behavior, aiming to further optimize particle manipulation in elasto-inertial microfluidics.

### Effect of Channel Geometry and Viscoelastic Properties on Nanoparticle Focusing

The focusing of nanoparticles in elasto-inertial microfluidics is strongly influenced by both channel geometry and fluid properties, such as the concentration of PEO. We systematically investigated these factors to optimize focusing behavior, using microchannel with different widths and viscoelastic fluids of varying PEO concentrations.

We used microchannels with widths of 5 µm and 10 µm, while keeping the channel height constant at 60 µm, resulting in aspect ratios of 12 and 6, respectively. As shown in Figure 3, fluorescence intensity profiles of 50 nm particles at PEO concentration of 1000 ppm and flow rates ranging from 0.5 µL/min to 3 µL/min indicate that particles are primarily focused at the channel center, independent of width and flow rate. However, a higher intensity signal at the channel center was observed in the narrower channel (*w = 5 µm*). This behavior likely results from stronger elastic forces in the smaller channel dimensions, where enhanced stress differences^40^ drive particles towards the minimum stress region at the channel center. Despite the consistent central focusing, the likelihood of unfocused particles increases with wider channels due to reduced elastic force. The focusing bandwidth (FWHM/w) is smaller across all flow rates for the narrower channel (*w = 5 µm*), with a significant increase in bandwidth observed at the highest flow rate of 3 µL/min in the wider channel (*w = 10 µm*) (see Figure S1). These trends are supported by numerical simulations (Figure S2), which reveal that channels with higher aspect ratios generate greater first normal stress differences (*N_1_*), intensifying elastic forces and improving focusing efficiency. Moreover, as expected, longer relaxation times (or higher Weissenberg numbers) correlate with increased elastic stresses and higher maximum *N_1_* (see Figure S2(a)). To the best of our knowledge, single-line particle focusing at the blockage ratios as low as 0.005 and 0.01 has not been achieved previously. We believe that the increased elasticity component and the utilization of high-aspect-ratio microchannels are the driving factors for nanoparticle focusing at such low blockage ratios.

**Figure 3.**
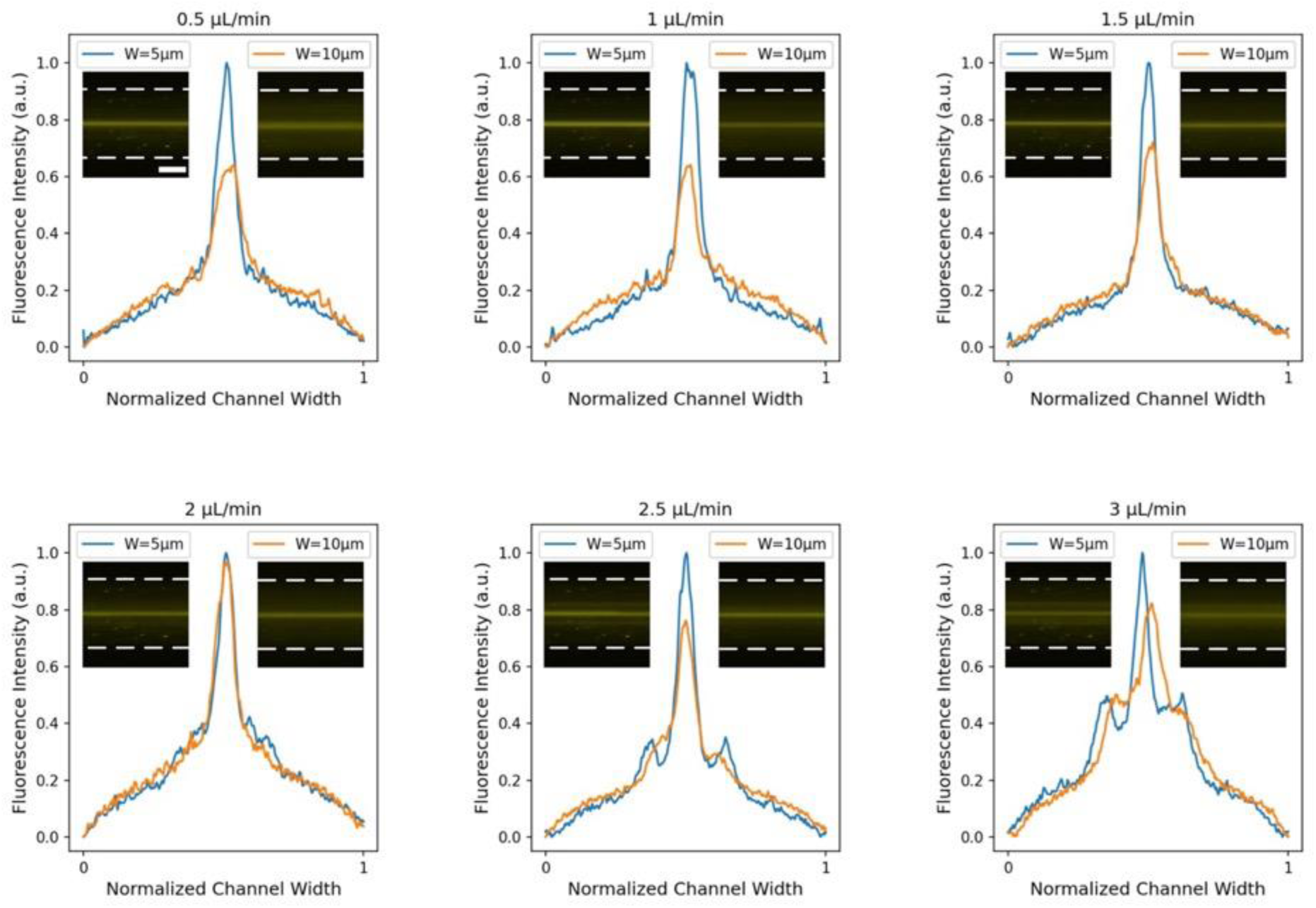
The effect of the channel width on particle focusing. Normalized intensity graphs of the cross-section comparing 50 nm particle focusing in microchannels with channel widths of 5 µm and 10 µm at a PEO concentration of 1000 ppm and flow rates ranging from 0.5 µL/min to 3 µL/min. The intensity profile highlights the effect of channel width on particle focusing quality. Scale bar: 50 µm.

The concentration of PEO significantly influences the viscoelastic fluid properties, which in turn govern the elasticity and viscosity components critical for elasto-inertial focusing. Table S1 details the rheological properties of the PEO solutions used in this study. Increasing PEO concentration enhances the relaxation time (λ) and fluid viscosity (μ), thereby strengthening the elastic component (*F_E_∼λ*)^41^ while reducing the Reynold number(*Re∼1/µ*).^42^

Figure 4 illustrates the impact of four different PEO concentrations (500 ppm, 1000 ppm, 2000 ppm, and 4000 ppm) on the focusing behavior of 25 nm and 100 nm particles in a microchannel with an aspect ratio (AR) of 12 (*h=60 µm, w=5 µm*) at a constant flow rate of 0.5 µL/min. Notably, the particles primarily focus at channel center even at the lowest PEO concentration of 500 ppm (Re=0.2, Wi=47.78, El=235.72), but some unfocused particles are observed. Increasing the concentration to 2000 ppm narrows the focusing bandwidth, eliminating unfocused particles around the center for the 100 nm particles. At 4000 ppm, focusing further improves for 100 nm particles, but no significant enhancement is observed for the 25 nm particles, which maintain their position at the center with slight distortions. These results suggest that increasing PEO concentration improved focusing quality for larger nanoparticles, but its effectiveness diminishes at smaller particle sizes due to competing effects between the different forces involved.

**Figure 4.**
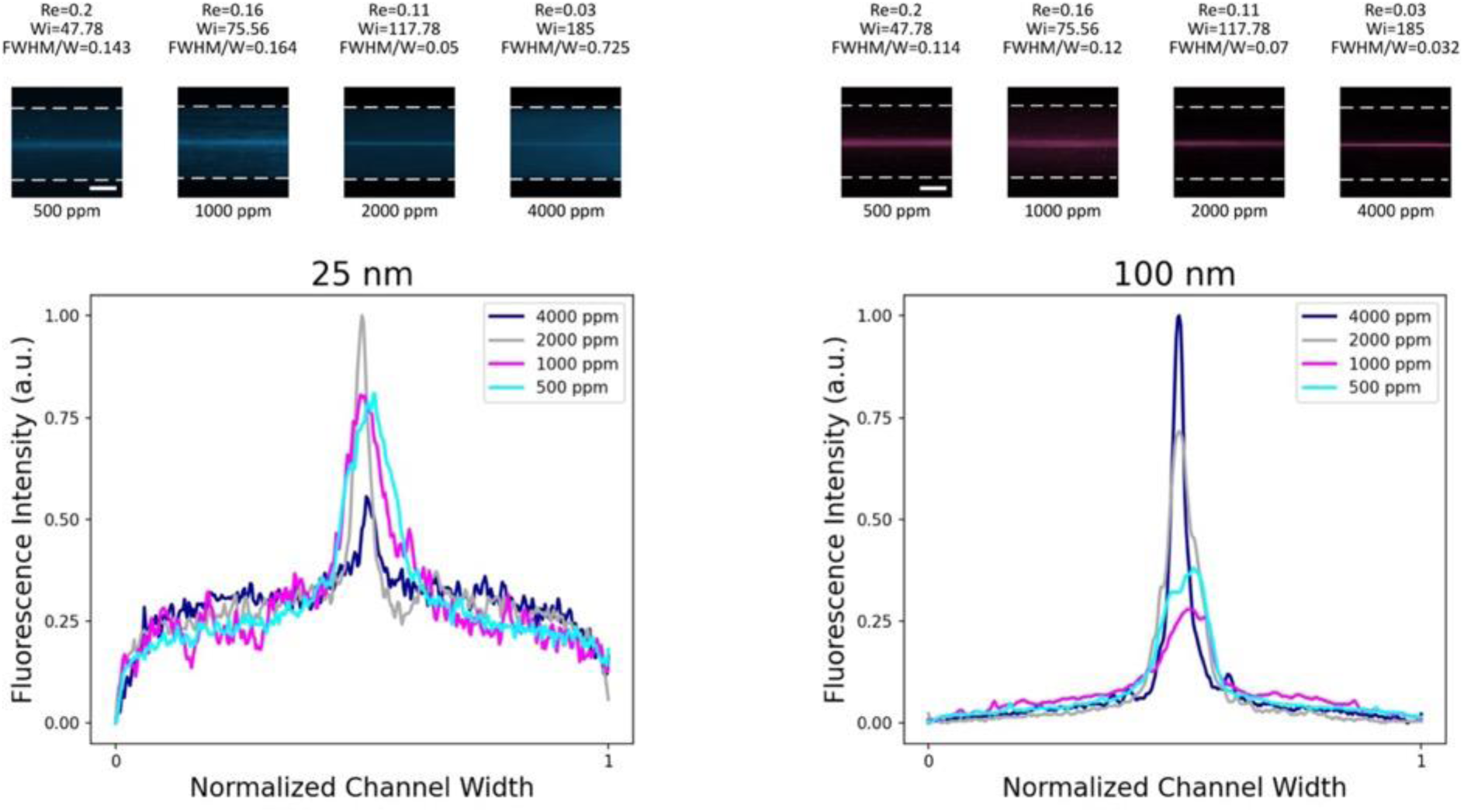
The effect of PEO concentration on particle focusing. Fluorescence images and normalized fluorescence intensity graphs of 25 nm and 100 nm particles at a constant flow rate of 0.5 µL/min. A higher PEO concentration improves focusing, especially for 100 nm particles. Scale bar: 50 µm.

The interplay between channel geometry and PEO concentration is crucial for achieving optimal nanoparticle focusing. Higher aspect ratio channels amplify elastic stresses, leading to sharper focusing, while increasing PEO concentrations enhances viscoelastic effects that further stabilize particle focusing and improve focusing efficiency. However, at higher concentrations, additional elasticity does not necessarily improve focusing for the smallest nanoparticles, indicating a need for balance between design parameters and fluid properties. These results highlight the importance of tailoring both geometric and fluidic parameters to the specific particle size and application requirements. With these insights into the effects of geometry and fluid properties, we now turn to the roles of particle size and flow rate in further refining the focusing behavior.

### Influence of Particle Size and Flow Rate on Nanoparticle Focusing

The performance of elasto-inertial microfluidics for nanoparticle focusing is strongly influenced by particle size and flow rate, which together determine the balance between elastic and inertial forces. To systematically investigate these effects, we studied six particle sizes (25 nm, 50 nm, 100 nm, 200 nm, 500 nm, and 1 µm) at a fixed PEO concentration of 2000 ppm in a microchannel with an aspect ratio (AR) of 12 (*h=60 µm, w=5 µm*) across a range of flow rates from 0.5 to 3 µL/min. Figures 5 and 6 summarize the experimental results, highlighting the interplay between these parameters.

**Figure 5.**
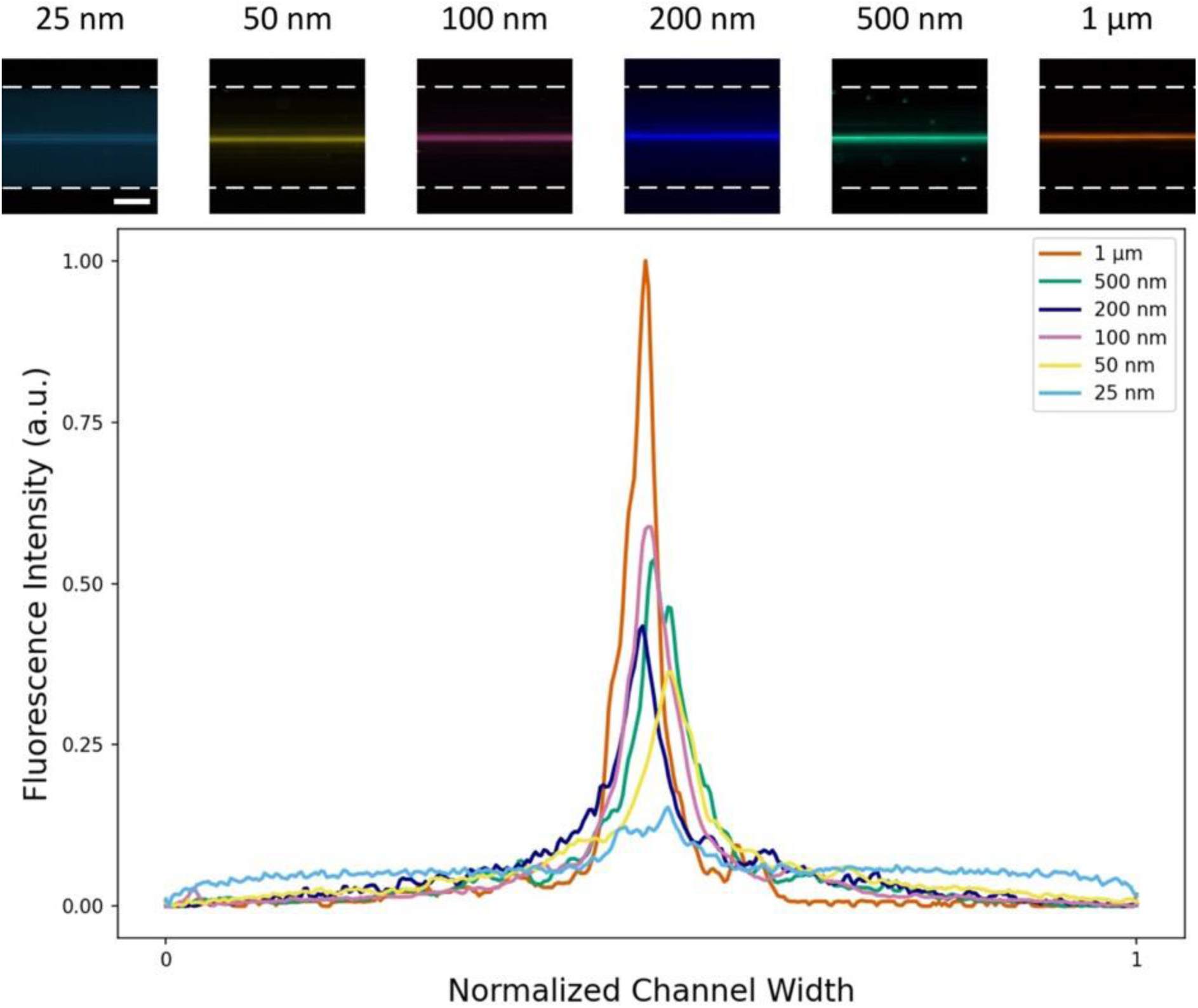
The effect of particle size. Fluorescence images and corresponding normalized fluorescence intensity graphs of particles ranging from 25 nm 1 µm at a PEO concentration of 2000 ppm and a flow rate of 1 µL/min. Larger particles exhibit higher intensity due to better focusing. Scale bar: 50 µm.

**Figure 6.**
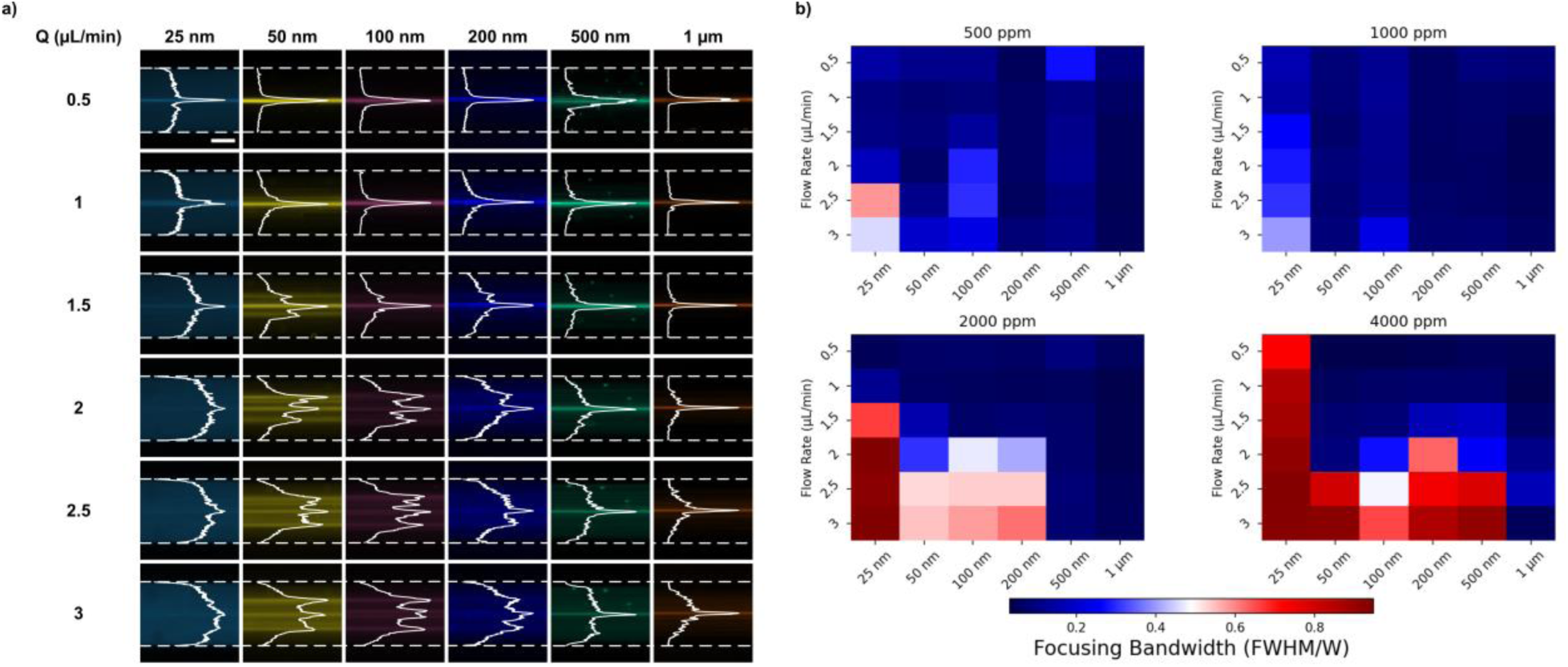
Effect of flow rate and PEO concentration. a) Fluorescence images of different particles (25 nm – 1 µm) are overlapped with corresponding cross-section intensity profiles to illustrate the focusing behavior at different flow rates. Scale bar: 50 µm. b) Heat map of focusing bandwidth for different particle sizes at varying flow rates and PEO concentrations. The data illustrate the interplay between fluid elasticity and flow rate.

At a fixed flow rate of 1 µL/min (Figure 5), all tested particles exhibit focusing at the channel center, with the sharpest fluorescence intensity profile observed for 1 µm particles. As particle size decreases, the fluorescence intensity at the center weakens, reflecting an increased proportion of unfocused particles. This trend suggests that larger particles experience stronger elastic forces (*F_E_∼a^3^, F_L_∼a^4^*),^43,44^ leading to a more confined and stable focusing profile, whereas smaller particles are more prone to Brownian motion and secondary flow effects. Notably, particles as small as 25 nm still demonstrate focusing behavior, albeit with broader intensity distribution compared to larger particles. These results confirm the feasibility of using elasto-inertial microfluidics for sub-100 nm particle focusing, overcoming previous limitations in label-free nanoparticle manipulation.

Beyond this fixed flow rate analysis, we explored the effect of increasing flow rate on nanoparticle focusing at the fixed PEO concentration of 2000 ppm (Figure 6). At lower flow rates (0.5 and 1 µL/min), all particle sizes predominantly focus at the channel center, with larger particles (500 nm and 1 µm) achieving the sharpest focusing profiles. Full width at half maximum (FWHM) analysis confirms that even the smallest particles (25 nm) exhibit focusing at the channel center, albeit with a broader intensity distribution. As flow rate increases to 1.5 µL/min, a transition occurs: smaller particles (50-200 nm) begin to develop additional focusing positions, while larger particles remain centered. At 2 µL/min and higher, three stable focusing positions emerge for the 50 nm and 100 nm particles—one at the channel center and two symmetrically positioned near the sides. The 200 nm particles also deviate from their single central position, instead showing a weaker three-position focusing pattern at 3 µL/min. In contrast, 500 nm and 1 µm particles maintain a single focused position at the channel center across all tested flow rates, though with decreasing fluorescence intensity at higher flow rates, suggesting a weaking of elastic forces as inertia grows dominant.

The loss of a distinct focusing at the channel center is expected in elasto-inertial focusing as the flow rate increases and inertia becomes stronger.^45^ These results align with force scaling predictions: elastic and lift forces scale differently with particle size (*F_E_∼a^3^, F_L_∼a^4^*), leading to stronger focusing of larger particles. However, the emergence of multiple focusing positions for smaller nanoparticles at higher flow rates suggests the influence of secondary flow effect in high-aspect-ratio microchannels. Previous studies have reported that secondary vortices can develop in straight channels at increased flow rates, displacing particles from the centerline due to transverse flow recirculation.^46,47^ Further research is required to fully characterize these effect in elasto-inertial focusing.

These findings suggest that for optimal nanoparticle focusing, lower flow rates (≤ 1 µL/min) are preferable, where elastic forces dominate over inertia. At these conditions, increasing PEO concentration enhances focusing quality by reinforcing viscoelastic effects. However, at higher flow rates, increasing inertia not only disrupts focusing but also introduces new stable focusing positions for smaller nanoparticles, requiring careful optimization for size-based separations.

In the following section, we further explore particle migration mechanisms for application in high-resolution nanoparticle separation, integrating numerical and experimental analyses.

### Particle Migration for Size-Based Separation

The ability to manipulate nanoparticles for size-based separation relies on a detailed understanding of their migration behavior in viscoelastic fluids. Our previous work demonstrated that prefocusing particles at the channel center is a crucial first step before achieving effective size-based separation.^39^ In this study, we extend this approach to nanoparticles and investigate both numerically and experimentally how elastic forces guide particle migration.

To elucidate the physical mechanisms underlying particle migration, we analyze the competition between elastic and inertial forces in a viscoelastic fluid. The shear gradient lift force pushes particles away from the centerline,^48^ while elastic forces counteract this by pulling the immersed particle toward the centerline for fluids which are not shear–thinning.^47,49–51^ As shown in Figure 7(a), numerical simulations of the first normal stress difference (*N*_1_ = τ_xx_ −τ_yy_) reveal that stress is highest near the channel walls and significantly lowers at the center, effectively creating a lateral thrust that drives particles to stable equilibrium positions.^50^ As a result, the gradient of *N*_1_ drives particles toward the channel center, where they encounter minimal elastic stress. This effect is particularly important in high-aspect-ratio channels, where it promotes single-line focusing.

**Figure 7.**
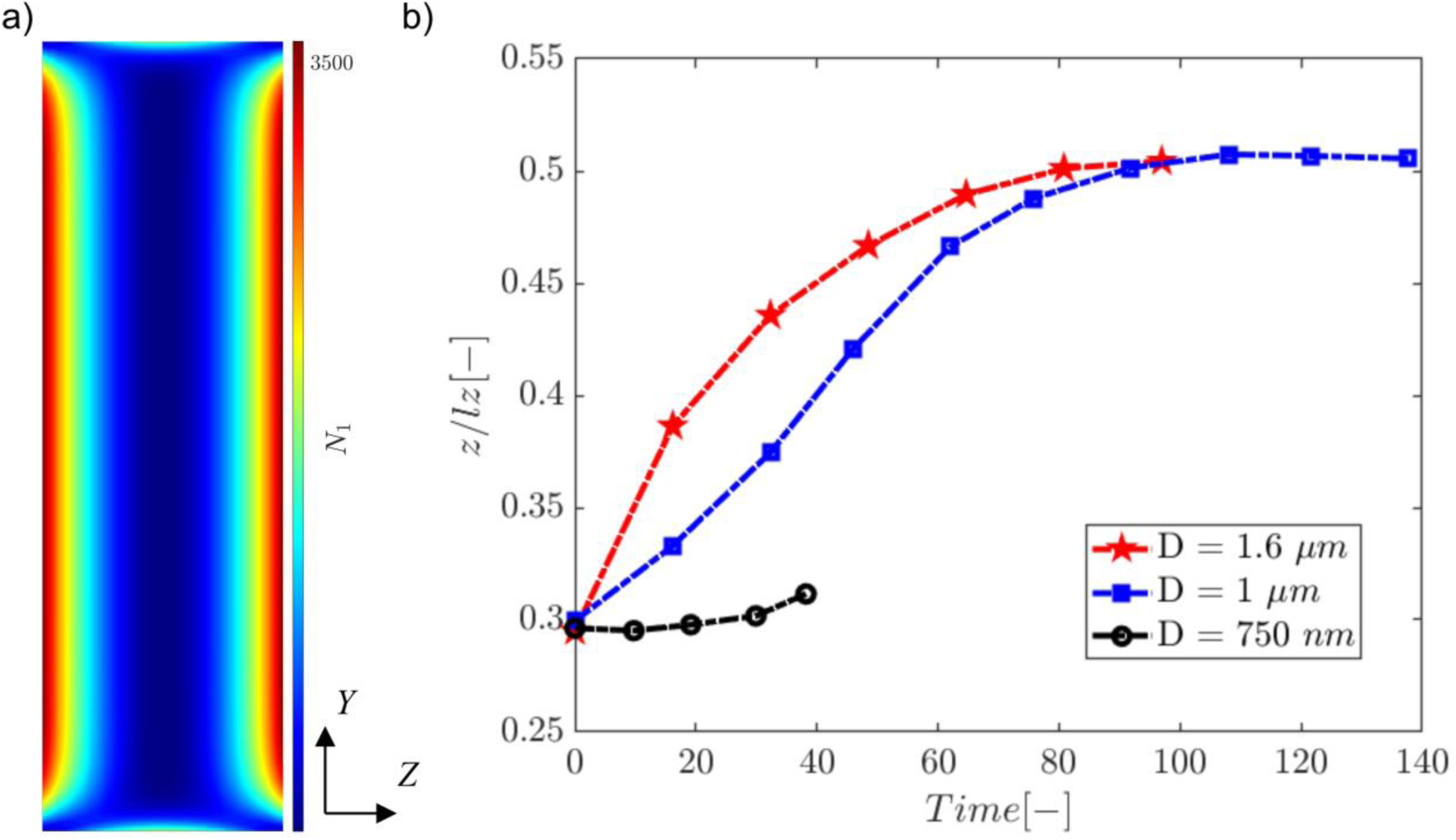
Numerical analysis of particle focusing and migration. a) The first normal stress difference (*N*_1_) across the channel cross–section. The particle reaches its equilibrium position at the center of the channel where *N*_1_=0. b) Particle spanwise position for AR=12 and the particles with diameters of 1.6 µm, 1 µm and 750 nm over time.

To quantify size-dependent migration, we performed numerical simulations for 1.6 µm, 1 µm, and 750 nm particles. As can be seen in Figure 7(b), all particles migrate toward the channel center (Z/l_Z_ = 0.5), but larger particles reach a steady-state position significantly faster than smaller ones. This is attributed to the scaling of elastic force, which increases with particle size (*F_E_∼a^3^*).

Consequently, 750 nm particles require more time to fully migrate, whereas 1 µm and 1.6 µm particles reach equilibrium more rapidly. The data clearly shows that the migration velocity of the largest particles is the highest. Moreover, migration velocity decreases as particles approach the centerline due to the progressively lower elastic stress gradients because of the lower local values of *N*_1_. The difference in migration velocity is attributed to elastic force arising from the imbalance in the distribution of *N*_1_ over the particle volume, *F*_E_ ∝ *a*^3^∇(*N*_1_), where “*a*” represents the radius of a spherical particle.^38,49^ Consequently, the larger particle experiences a more pronounced elastic force, which accelerates its migration toward the centerline more than the smaller particles.

It is important to highlight that while particle migration along the *Z*-axis is consistently substantial, cross-stream motions along the *Y*-axis are less pronounced. This can be again explained by the distribution of the first normal stress difference (*N*_1_) within the rectangular channel. In other words, while migration along the primary axis (z-direction) is well-defined, cross-stream migration along the lateral axis (y-direction) remains limited, as the lowest *N*_1_ values form a vertically extended region, allowing for consistent particle alignment along this plane.

To validate these numerical predictions, we conducted experimental studies tracking the migration of 500 nm and 1 µm particles in a two-stage microfluidic device (Figure 8). In the first section, particles were prefocused at the channel center, after which the channel split into two branches, enabling observation of migration back toward the centerline. The results confirm that 1 µm particles migrate significantly faster than 500 nm particles, in agreement with numerical predictions. Figure 8(b) shows the intensity profile for the 1 µm particles and 500 nm particles where the 1 µm particles migrate and reach the centerline faster while the 500 nm particles are lagging behind and exhibit broader spatial distribution, increasing overlap with 1 µm particles, which could pose challenges for high-purity separation.

**Figure 8.**
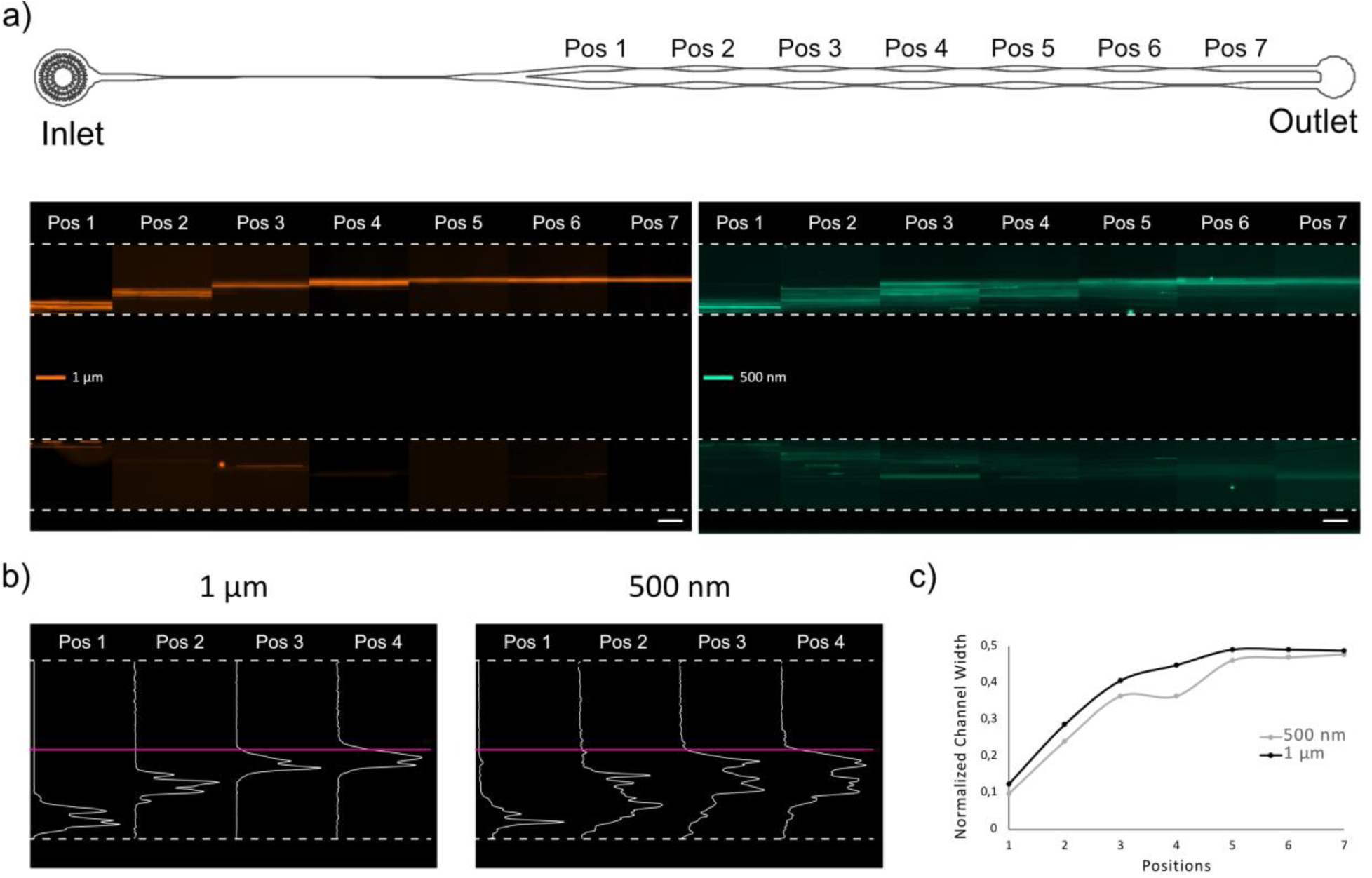
Particle pre-focusing and migration for separation. a) Schematic of a microfluidic chip with the pre-focusing and migration sections, where fluorescence images are taken to observe migration of 500 nm and 1 µm particles. Scale bar: 50 µm b) The fluorescence intensity profiles of both 500 nm and 1 µm particles. The pink line indicates the center of the channel c) Trajectories of both particles based on average intensity points. The larger (1 µm) particles migrate faster to the center.

However, further optimizations are needed to achieve complete size-based separation since 500 nm particles are spread wider while migrating towards the channel center, overlapping with 1 µm particles. Under similar conditions, we successfully separate 2 µm and 3 µm particles, indicating that separation resolution is size-dependent (see Figure S3), with smaller nanoparticles requiring additional optimizations.

These findings demonstrate that elasto-inertial migration can effectively enrich nanoparticles, but further refinements are needed to achieve complete separation at the sub-micron scale. At this size range, Brownian motion and reduced elastic force gradients introduce challenges that must be addressed by optimizing channel geometry, flow conditions, and viscoelastic properties. Despite these complexities, the observed migration behavior paves the way for high-resolution nanoparticle sorting using elasto-inertial microfluidics. Future efforts will focus on multi-stage separation strategies and further tuning of viscoelastic stress distributions to enhance separation fidelity.

### Focusing of Biological Nanoparticles

To demonstrate the potential of elasto-inertial microfluidics in high-aspect-ratio microchannels for biomedical applications, we investigated the focusing behavior of biologically relevant nanoparticles. We conducted experiments using high-density lipoproteins (HDL, 10 nm), low-density lipoproteins (LDL, 25 nm), liposomes (90 nm), and extracellular vesicles (EVs, 100 nm) in a viscoelastic fluid at 1000 ppm PEO (detailed size distribution of biological particles are provided in Figure S4).

Our results, presented in Figure 9, reveal that biological particles exhibit central focusing within the microchannel, consistent with the behavior observed for polystyrene nanoparticles. The smallest particles (HDL, 10 nm) exhibited the highest background signal, likely due to their small size and increased diffusion. However, LDL (25 nm), liposomes (90 nm), and EVs (100 nm) were fully focused at the channel center, demonstrating the ability of elasto-inertial forces to counteract Brownian motion and drive nanoscale bioparticles to a single equilibrium position.

**Figure 9.**
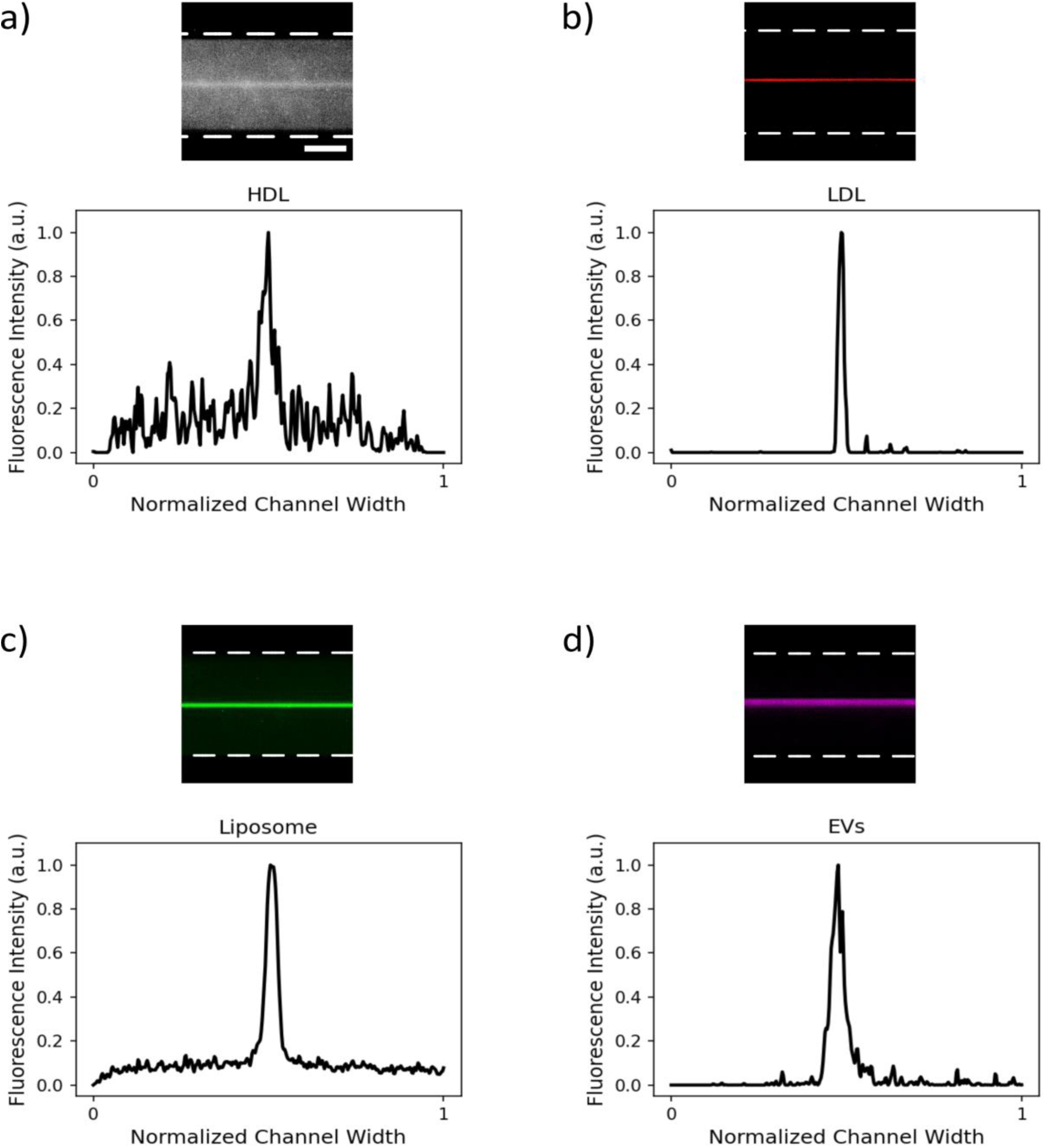
Focusing of biological nanoparticles (HDL, LDL, liposomes, EVs) in high-aspect-ratio microchannels at a PEO concentration of 1000 ppm and a flow rate of 1 µL/min. a) HDL (10 nm), b) LDL (25 nm), c) Liposome (90 nm), d) EVs (100 nm). Scale bar: 50 µm. Results demonstrate the applicability of elasto-inertial focusing for biological samples.

These findings establish the feasibility of elasto-inertial focusing for the manipulation of biological nanoparticles, which is crucial for applications requiring particle enrichment, high-purity isolation, and potential size-based separation. Unlike conventional methods such as size-exclusion chromatography, ultracentrifugation, and asymmetrical flow field-flow fractionation (AF4), which often require multiple processing steps and specialized equipment, elasto-inertial microfluidics provides a label-free, single-step approach with minimal sample preparation. Additionally, compared to microfluidics based active methods such as acoustofluidics and dielectrophoresis, which rely on external fields and complex device architectures,^52–54^ the method presented here enables high-throughput nanoparticle manipulation with a simple, single-inlet design.

Previous studies have explored viscoelastic microfluidics for extracellular vesicle and lipoprotein enrichment, but these methods have been limited in their ability to precisely focus particles below 50 nm in sheathless conditions. The present work extends the lower focusing limit to 10 nm, demonstrating that high-aspect-ratio channels combined with optimized viscoelastic conditions provide a scalable platform for sub-100 nm particle manipulation. Beyond focusing, our findings lay the groundwork for future developments in size-based separation of nanoparticles (<500 nm) using elasto-inertial forces.

## CONCLUSION

This study establishes elasto-inertial focusing of nanoparticles as small as 25 nm in a sheathless flow, high-aspect-ratio channels. While elasto-inertial microfluidics has previously demonstrated advantages for microscale particle manipulation, its application to nanoscale particles has been significantly limited. While previous research demonstrated focusing down to 200 nm in straight microchannels, we extend this limit to 10 nm, marking a significant leap in elasto-inertial microfluidics by achieving nanoparticle focusing in high-aspect-ratio microchannels.

Numerical and experimental analyses together provide a deeper understanding of particle migration dynamics in elasticity-dominated flows (0.03<Re<1.22, 11<Wi<1110, 63<El<6000), reinforcing the robustness of this approach and its potential for broad applications. Our systematic study confirms the feasibility of elasto-inertial focusing for biologically relevant nanoparticles, including HDL (10 nm), LDL (25 nm), lipoproteins (90 – 140 nm) and EVs (100 nm), under optimized flow conditions. The ability to focus biologically relevant nanoparticles, including those used in drug delivery (e.g., liposomes for mRNA vaccines), opens new avenues for high-resolution fractionation of biomolecular carries, with implications for both diagnostics and therapeutic applications. We envision that this approach will pave the way for next-generation microfluidic systems capable of high-throughput particle separation, bioparticle enrichment, and precision medicine applications. Future work will explore integration with downstream analytical techniques and the development of parallelized systems for enhanced throughput.

## MATERIALS AND METHODS

### Device Fabrication

Microfluidic devices were designed with the AutoCAD software (Autodesk). The master mold to fabricate the PDMS (polydimethylsiloxane) chips was prepared with SUEX (dry SU-8 film) on a silicon wafer through photolithography process.^55^ The PDMS base (SYLGARD 184) was mixed with the curing agent at the mixing ratio of 10:1. The mixture was poured onto the master mold. Degassed in a desiccator and baked at 65 ℃ for 6 hours for curing. The cured PDMS was peeled off, and inlet and outlet holes were punched. The prepared PDMS chip and microscope glass slide were bonded using oxygen plasma activation. The final device was post-baked at 120 ℃ for 15 minutes for better sealing.

### Design of Microfluidic Channels

Two different high aspect ratio microfluidic channels were designed to study the nanoparticle focusing. Both microfluidic channels contain 1.5 mm long straight channel with width 20 µm followed by 3 mm long straight channel, which is then followed by another 1.5 mm long straight channel with width 20 µm. The only difference between the wo microfluidic channels is the width of the 3 mm long straight channel. First design has 5 µm width (AR=12) while the second design has 10 µm width (AR=6). Two different widths are therefore used to investigate the effect of the channel geometry. The channel height was kept constant at 60 µm for all the experiments. Both channels have an expansion area (width=150 µm) before the outlet to observe the particles in high-resolution. (See Figure 1 for the channel design). The data used and analyzed in this study were recorded in these expansion areas.

### Sample Preparation

In this study, PEO (Polyethylene Oxide, M_w_=2×10^6^ g/mol), elasticity enhancer, was used for the preparation of viscoelastic fluids. The PEO powder was dissolved in deionized water at four different concentrations (500 ppm, 1000 ppm, 2000 ppm and 4000 ppm) (See Table S1 for the rheological properties of the fluids and see Table S2 for the dimensionless numbers). The fluorescent polystyrene (PS) particles (Fluoro-Max, ThermoFisher Scientific) with diameters of 25 nm, 50 nm, 100 nm, 200 nm, 500 nm and 1 µm were suspended in the prepared viscoelastic fluids prior to the flow experiments (3 µL of particles at the concentration of 1% solids were suspended at 3 mL of PEO).

### Numerical Analysis

We perform three-dimensional direct numerical simulations to investigate the cross-streamline migration of particles suspended in viscoelastic (VE) fluids within a relatively high aspect ratio straight microchannel. These simulations aim to further confirm and explain the experimental observations. We employ our in-house code utilizing a direct forcing immersed boundary method (IBM) to simulate the particles as moving Lagrangian grids, while the carrier fluid is discretized within a stationary Eulerian frame, in which the Navier-Stokes and the viscoelastic constitutive equations are discretized using finite differences.^56^ The suspending fluid motion is governed by the incompressibility constraint and conservation of momentum as follows:

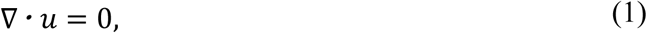

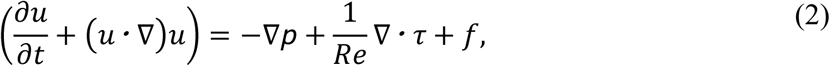

Here, *u* is the fluid velocity, *p* is the pressure field, τ is the total deviatoric stress tensor, and Re is Reynolds number. The extra term f on the right-hand side of Equation (2) is the immersed boundary force field representing the particle-fluid interaction; details of immersed boundary method can be found in the work of Breugem.^57^ The total deviatoric stress tensor, τ, is composed of contributions from the solvent (Newtonian fluid) and polymer parts as τ = τ^*p*^ + τ^*s*^. The solvent stress tensor is defined as τ^*s*^ = β_*s*_ (∇*u* + ∇*u*^T^), where β_*s*_ = μ_*s*_/μ is the ratio of the solvent viscosity to the total viscosity. In addition to the equations mentioned earlier, a constitutive equation should be employed to model the evolution of the non-Newtonian contribution (τ^*p*^) of the VE material. We implement the Oldroyd-B model^58^ to consider the viscoelasticity of the material.

### Preparation of Biological Nanoparticles

Lipoproteins were isolated from the blood plasma of apparently healthy donors, obtained from Blood transfusion station of Karolinska Hospital. The density of 3 ml of blood plasma was adjusted to 1.021 g/mL using KBr powder. The KBr buffers of different densities (1.006 g/ml, 1.019g/ml, 1.063 g/mL, 1.24 g/ml) and blood plasma were layered in a gradient, transitioning from denser to less dense solutions in 14 mL tubes (14 x 95 mm, Open-Top Thinwall Ultra-Clear Tubes, Backman Coulter, USA). The tubes were ultracentrifuged for 48 hours at 37000 rpm at 15°C (Optima-XE Ultracentrifuge, SW 40 Ti Swinging-Bucket Rotor, Backman Coulter, USA). Post-centrifugation, LP fractions containing separately Very Low-Density Lipoproteins (VLDL), Low Density Lipoproteins (LDL) and High-Density Lipoproteins (HDL) were carefully removed using a medical needle and syringe avoiding the blending of different classes of LPs. Furthermore, VLDL, LDL and HDL were concentrated using Ultra Centrifugal Filter, 30 kDa MWCO (Amicon). Protein concentration was measured using Bradford assay (Biorad) at NanoDrop Microvolume Spectrophotometer (ThermoFisher). The sizes of lipoproteins were confirmed using Dynamic Light Scattering at Zeta-sizer (Malvern). HDL and LDL were labeled with TopFluor-Cholesterol (Avanti Polar Lipids). Finally, 50 µL of HDL and LDL particles at the concentration of 0.5 mg/mL were diluted to 1 mL with PEO prior the flow experiments.

For production of extracellular vesicles (EVs), immortalized fibroblasts (BJ-5ta cell line, ATCC CRL-4001) were cultured in a 4:1 mixture of Dulbecco’s medium supplemented with 4 mM L-glutamine, glucose (4.5 g/L) and sodium bicarbonate (1.5 g/liter) and Medium 199 supplemented with Hygromycin B (0.01 mg/mL) (ThermoFisher Scientific) and 10% fetal bovine serum (Invitrogen). Cells were cultured at 37°C and 5% CO_2_ in a humidified atmosphere and regularly tested for the presence of mycoplasma. For EV harvesting, media was changed to OptiMem (Invitrogen) 48 hours before harvest of conditioned media (CM) as described before.^59^ Collected CM was directly subjected to a low-speed centrifugation step at 500 x g for 5 min followed by a 2000 x g spin for 10 min to remove larger particles and cell debris. Precleared CM was subsequently filtered through 0.22 μm bottle top vacuum filters (Corning, cellulose acetate, low protein binding) to remove any larger particles. EVs were then prepared by tangential flow filtration (TFF) as described before.^60^ In brief, precleared CM was concentrated via TFF by using the KR2i TFF system (Spectrum Labs) equipped with modified polyethersulfone hollow fiber filters with 300 kDa membrane pore size (MidiKros, 370 cm^2^ surface area, Spectrum Labs) at a flow rate of 100 mL/min (transmembrane pressure at 3.0 psi and shear rate at 3700 sec^−1^) as described previously. Amicon Ultra-0.5 10 kDa MWCO spin-filters (Millipore) were used to concentrate the sample to a final volume of 100 μL. Final EV samples were stored at −80°C in PBS-HAT [PBS supplemented with HEPES, human serum albumin and D-(+)-Trehalose dihydrate] until usage.^61^

Synthetic Liposomes were prepared using the lipid 1,2-Dioleoyl-sn-glycero-3-phosphocholine (DOPC). DOPC was dissolved in 2 ml of chloroform at 0.25 mg/ml in the glass vial. Chloroform was then evaporated under the nitrogen flow. The lipid residue was hydrated using the buffer (150 mM of NaCl, 10 mM of HEPES) and vortexed harshly until the solution became turbid whereas the lipid residue disappeared from the wall of the glass vial. Then, the liquid was transferred to the 15 ml falcon tube and sonicated at power 3, duty cycle 40%, for 10 min using a Branson Sonifier 250. The second aliquot, instead of sonication was extruded 21 times using the mini extruder (Avanti) with filter size of 100 nm. The size was verified using DLS (Malvern) (see Figures S3).

### Experimental Setup and Analysis

The microfluidic experiments were performed with a mid-pressure pump (neMESYS CETONI GmbH) using a 3 mL steel syringe. The data acquisition was accomplished using an inverted microscope (Nikon Eclipse TI) with a sCMOS camera (Andor Zyla) and LED lightning system (Lumenor Spectra X LED). Micro Manager software was used to control the microscope and record the images. The recorded images were processed by ImageJ software.

## AUTHOR INFORMATION

### Author Contributions

S.T.: conceptualization, validation, formal analysis, investigation, writing-original draft, writing-review and editing. J.C.: conceptualization, resources, writing-review and writing. S.H.: conceptualization, formal analysis, investigation, writing-review and editing. T.S.: conceptualization, resources, writing-review and editing. M.C.: formal analysis, writing-review and editing. G.M.: conceptualization, writing-review and editing, supervision. A.G.: resources, writing-review and editing. S.A.: resources, supervision. L.B.: writing-review and editing, supervision. O.T.: writing-review and editing, supervision. E.S.: conceptualization, resources, writing-review and editing, supervision. A.R.: conceptualization, writing-review and editing, supervision, project administration, funding acquisition.

## ACKNOWLEDGMENT

This research has received funding from the European Union’s Framework Programme for Research and Innovation Horizon 2020 under the Marie Skłodowska-Curie Grant Agreement No. 860775, the European Union’s Horizon 2020 research and innovation program under the Marie Skłodowska-Curie grant agreement No. 955605, the Horizon Europe research and innovation program under grant agreement No. 101057596, the Swedish Research Council (VR 2021-05861). The authors acknowledge the computational resources provided by the National Academic Infrastructure for Supercomputing in Sweden (NAISS). We also gratefully acknowledge the support of the European Research Council through Starting Grant MUCUS (Grant No. ERC-StG-2019-852529).

## REFERENCES

(1) Hettiarachchi, S.; Cha, H.; Ouyang, L.; Mudugamuwa, A.; An, H.; Kijanka, G.; Kashaninejad, N.; Nguyen, N.-T.; Zhang, J. Recent Microfluidic Advances in Submicron to Nanoparticle Manipulation and Separation. Lab. Chip 2023, 23 (5), 982–1010. 10.1039/D2LC00793B.

(2) Lin, M.; Li, M.; Zheng, H.; Sun, H.; Zhang, J. Lipoprotein Proteome Profile: Novel Insight into Hyperlipidemia. Clin. Transl. Med. 2021, 11 (4), e361. 10.1002/ctm2.361.

(3) Martín-Peña, A.; Tansey, M. G. Alzheimer’s Risk Gene Works the Gut–Brain Axis.

(4) Blanchard, J. W.; Akay, L. A.; Davila-Velderrain, J.; Von Maydell, D.; Mathys, H.; Davidson, S. M.; Effenberger, A.; Chen, C.-Y.; Maner-Smith, K.; Hajjar, I.; Ortlund, E. A.; Bula, M.; Agbas, E.; Ng, A.; Jiang, X.; Kahn, M.; Blanco-Duque, C.; Lavoie, N.; Liu, L.; Reyes, R.; Lin, Y.-T.; Ko, T.; R’Bibo, L.; Ralvenius, W. T.; Bennett, D. A.; Cam, H. P.; Kellis, M.; Tsai, L.-H. APOE4 Impairs Myelination via Cholesterol Dysregulation in Oligodendrocytes. Nature 2022, 611 (7937), 769–779. 10.1038/s41586-022-05439-w.

(5) Logozzi, M.; De Milito, A.; Lugini, L.; Borghi, M.; Calabrò, L.; Spada, M.; Perdicchio, M.; Marino, M. L.; Federici, C.; Iessi, E.; Brambilla, D.; Venturi, G.; Lozupone, F.; Santinami, M.; Huber, V.; Maio, M.; Rivoltini, L.; Fais, S. High Levels of Exosomes Expressing CD63 and Caveolin-1 in Plasma of Melanoma Patients. PLoS ONE 2009, 4 (4), e5219. 10.1371/journal.pone.0005219.

(6) Robertson, J. D.; Rizzello, L.; Avila-Olias, M.; Gaitzsch, J.; Contini, C.; Magoń, M. S.; Renshaw, S. A.; Battaglia, G. Purification of Nanoparticles by Size and Shape. Sci. Rep. 2016, 6 (1), 27494. 10.1038/srep27494.

(7) Quattrini, F.; Berrecoso, G.; Crecente-Campo, J.; Alonso, M. J. Asymmetric Flow Field-Flow Fractionation as a Multifunctional Technique for the Characterization of Polymeric Nanocarriers. Drug Deliv. Transl. Res. 2021, 11 (2), 373–395. 10.1007/s13346-021-00918-5.

(8) Xu, X.; Cölfen, H. Ultracentrifugation Techniques for the Ordering of Nanoparticles. Nanomaterials 2021, 11 (2), 333. 10.3390/nano11020333.

(9) Bostan, N.; Ilyas, N.; Akhtar, N.; Mehmood, S.; Saman, R. U.; Sayyed, R. Z.; Shatid, A. A.; Alfaifi, M. Y.; Elbehairi, S. E. I.; Pandiaraj, S. Toxicity Assessment of Microplastic (MPs); a Threat to the Ecosystem. Environ. Res. 2023, 234, 116523. 10.1016/j.envres.2023.116523.

(10) Lai, H.; Liu, X.; Qu, M. Nanoplastics and Human Health: Hazard Identification and Biointerface. Nanomaterials 2022, 12 (8), 1298. 10.3390/nano12081298.

(11) Preetam, S.; Nahak, B. K.; Patra, S.; Toncu, D. C.; Park, S.; Syväjärvi, M.; Orive, G.; Tiwari, A. Emergence of Microfluidics for next Generation Biomedical Devices. Biosens. Bioelectron. X 2022, 10, 100106. 10.1016/j.biosx.2022.100106.

(12) Liu, Y.; Jiang, X. Why Microfluidics? Merits and Trends in Chemical Synthesis. Lab. Chip 2017, 17 (23), 3960–3978. 10.1039/C7LC00627F.

(13) Zhu, X.; Wang, K.; Yan, H.; Liu, C.; Zhu, X.; Chen, B. Microfluidics as an Emerging Platform for Exploring Soil Environmental Processes: A Critical Review. Environ. Sci. Technol. 2022, 56 (2), 711–731. 10.1021/acs.est.1c03899.

(14) Whitesides, G. M. The Origins and the Future of Microfluidics. Nature 2006, 442 (7101), 368–373. 10.1038/nature05058.

(15) Dimaki, M.; Olsen, M. H.; Rozlosnik, N.; Svendsen, W. E. Sub–100 Nm Nanoparticle Upconcentration in Flow by Dielectrophoretic Forces. Micromachines 2022, 13 (6), 866. 10.3390/mi13060866.

(16) Zhang, Y.; Zhou, A.; Chen, S.; Lum, G. Z.; Zhang, X. A Perspective on Magnetic Microfluidics: Towards an Intelligent Future. Biomicrofluidics 2022, 16 (1), 011301. 10.1063/5.0079464.

(17) Wu, M.; Ozcelik, A.; Rufo, J.; Wang, Z.; Fang, R.; Jun Huang, T. Acoustofluidic Separation of Cells and Particles. Microsyst. Nanoeng. 2019, 5 (1), 32. 10.1038/s41378-019-0064-3.

(18) Costa, M.; Hammarström, B.; Van Der Geer, L.; Tanriverdi, S.; Joensson, H. N.; Wiklund, M.; Russom, A. EchoGrid: High-Throughput Acoustic Trapping for Enrichment of Environmental Microplastics. Anal. Chem. 2024, 96 (23), 9493–9502. 10.1021/acs.analchem.4c00933.

(19) Martel, J. M.; Toner, M. Inertial Focusing in Microfluidics. Annu. Rev. Biomed. Eng. 2014, 16 (1), 371–396. 10.1146/annurev-bioeng-121813-120704.

(20) Zhou, J.; Papautsky, I. Viscoelastic Microfluidics: Progress and Challenges. Microsyst. Nanoeng. 2020, 6 (1), 113. 10.1038/s41378-020-00218-x.

(21) Bhagat, A. A. S.; Kuntaegowdanahalli, S. S.; Papautsky, I. Enhanced Particle Filtration in Straight Microchannels Using Shear-Modulated Inertial Migration. Phys. Fluids 2008, 20 (10), 101702. 10.1063/1.2998844.

(22) Segré, G.; Silberberg, A. Radial Particle Displacements in Poiseuille Flow of Suspensions. Nature 1961, 189 (4760), 209–210. 10.1038/189209a0.

(23) Cruz, J.; Hjort, K. The Upper Limit and Lift Force within Inertial Focusing in High Aspect Ratio Curved Microfluidics. Sci. Rep. 2021, 11 (1), 6473. 10.1038/s41598-021-85910-2.

(24) Cruz, J.; Hjort, K.; Hjort, K. Stable 3D Inertial Focusing by High Aspect Ratio Curved Microfluidics. J. Micromechanics Microengineering 2021, 31 (1), 015008. 10.1088/1361-6439/abcae7.

(25) Cruz, J.; Hjort, K. High-Resolution Particle Separation by Inertial Focusing in High Aspect Ratio Curved Microfluidics. Sci. Rep. 2021, 11 (1), 13959. 10.1038/s41598-021-93177-w.

(26) Leshansky, A. M.; Bransky, A.; Korin, N.; Dinnar, U. Tunable Nonlinear Viscoelastic “Focusing” in a Microfluidic Device. Phys. Rev. Lett. 2007, 98 (23), 234501. 10.1103/PhysRevLett.98.234501.

(27) Liu, C.; Xue, C.; Chen, X.; Shan, L.; Tian, Y.; Hu, G. Size-Based Separation of Particles and Cells Utilizing Viscoelastic Effects in Straight Microchannels. Anal. Chem. 2015, 87 (12), 6041–6048. 10.1021/acs.analchem.5b00516.

(28) De Santo, I.; D’Avino, G.; Romeo, G.; Greco, F.; Netti, P. A.; Maffettone, P. L. Microfluidic Lagrangian Trap for Brownian Particles: Three-Dimensional Focusing down to the Nanoscale. Phys. Rev. Appl. 2014, 2 (6), 064001. 10.1103/PhysRevApplied.2.064001.

(29) Liu, C.; Ding, B.; Xue, C.; Tian, Y.; Hu, G.; Sun, J. Sheathless Focusing and Separation of Diverse Nanoparticles in Viscoelastic Solutions with Minimized Shear Thinning. Anal. Chem. 2016, 88 (24), 12547–12553. 10.1021/acs.analchem.6b04564.

(30) Young Kim, J.; Won Ahn, S.; Sik Lee, S.; Min Kim, J. Lateral Migration and Focusing of Colloidal Particles and DNA Molecules under Viscoelastic Flow. Lab. Chip 2012, 12 (16), 2807. 10.1039/c2lc40147a.

(31) Li, D.; Xuan, X. Fluid Rheological Effects on Particle Migration in a Straight Rectangular Microchannel. Microfluid. Nanofluidics 2018, 22 (4), 49. 10.1007/s10404-018-2070-4.

(32) D’Avino, G.; Maffettone, P. L. Particle Dynamics in Viscoelastic Liquids. J. Non-Newton. Fluid Mech. 2015, 215, 80–104. 10.1016/j.jnnfm.2014.09.014.

(33) Zhou, J.; Papautsky, I. Fundamentals of Inertial Focusing in Microchannels. Lab. Chip 2013, 13 (6), 1121. 10.1039/c2lc41248a.

(34) Lim, H.; Nam, J.; Shin, S. Lateral Migration of Particles Suspended in Viscoelastic Fluids in a Microchannel Flow. Microfluid. Nanofluidics 2014, 17 (4), 683–692. 10.1007/s10404-014-1353-7.

(35) Li, D.; Lu, X.; Xuan, X. Viscoelastic Separation of Particles by Size in Straight Rectangular Microchannels: A Parametric Study for a Refined Understanding. Anal. Chem. 2016, 88 (24), 12303–12309. 10.1021/acs.analchem.6b03501.

(36) Zhou, J.; Giridhar, P. V.; Kasper, S.; Papautsky, I. Modulation of Aspect Ratio for Complete Separation in an Inertial Microfluidic Channel. Lab. Chip 2013, 13 (10), 1919. 10.1039/c3lc50101a.

(37) Di Carlo, D. Inertial Microfluidics. Lab. Chip 2009, 9 (21), 3038. 10.1039/b912547g.

(38) Tehrani, M. A. An Experimental Study of Particle Migration in Pipe Flow of Viscoelastic Fluids. J. Rheol. 1996, 40 (6), 1057–1077. 10.1122/1.550773.

(39) Tanriverdi, S.; Cruz, J.; Habibi, S.; Amini, K.; Costa, M.; Lundell, F.; Mårtensson, G.; Brandt, L.; Tammisola, O.; Russom, A. Elasto-Inertial Focusing and Particle Migration in High Aspect Ratio Microchannels for High-Throughput Separation. Microsyst. Nanoeng. 2024, 10 (1), 87. 10.1038/s41378-024-00724-2.

(40) Lu, X.; Zhu, L.; Hua, R.; Xuan, X. Continuous Sheath-Free Separation of Particles by Shape in Viscoelastic Fluids. Appl. Phys. Lett. 2015, 107 (26), 264102. 10.1063/1.4939267.

(41) Kumar, T.; Ramachandraiah, H.; Iyengar, S. N.; Banerjee, I.; Mårtensson, G.; Russom, A. High Throughput Viscoelastic Particle Focusing and Separation in Spiral Microchannels. Sci. Rep. 2021, 11 (1), 8467. 10.1038/s41598-021-88047-4.

(42) Rott, N. Note on the History of the Reynolds Number.

(43) Zhou, Y.; Ma, Z.; Ai, Y. Dynamically Tunable Elasto-Inertial Particle Focusing and Sorting in Microfluidics. Lab. Chip 2020, 20 (3), 568–581. 10.1039/C9LC01071H.

(44) Xiang, N.; Zhang, X.; Dai, Q.; Cheng, J.; Chen, K.; Ni, Z. Fundamentals of Elasto-Inertial Particle Focusing in Curved Microfluidic Channels. Lab. Chip 2016, 16 (14), 2626–2635. 10.1039/C6LC00376A.

(45) Naderi, M. M.; Barilla, L.; Zhou, J.; Papautsky, I.; Peng, Z. Elasto-Inertial Focusing Mechanisms of Particles in Shear-Thinning Viscoelastic Fluid in Rectangular Microchannels. Micromachines 2022, 13 (12), 2131. 10.3390/mi13122131.

(46) Xue, S.-C.; Phan-Thien, N.; Tanner, R. I. Numerical Study of Secondary Flows of Viscoelastic Fluid in Straight Pipes by an Implicit Finite Volume Method. J. Non-Newton. Fluid Mech. 1995, 59 (2–3), 191–213. 10.1016/0377-0257(95)01365-3.

(47) Li, G.; McKinley, G. H.; Ardekani, A. M. Dynamics of Particle Migration in Channel Flow of Viscoelastic Fluids. J. Fluid Mech. 2015, 785, 486–505. 10.1017/jfm.2015.619.

(48) Asmolov, E. S. The Inertial Lift on a Spherical Particle in a Plane Poiseuille Flow at Large Channel Reynolds Number. J. Fluid Mech. 1999, 381, 63–87. 10.1017/S0022112098003474.

(49) Karimi, A.; Yazdi, S.; Ardekani, A. M. Hydrodynamic Mechanisms of Cell and Particle Trapping in Microfluidics. Biomicrofluidics 2013, 7 (2), 021501. 10.1063/1.4799787.

(50) Ho, B. P.; Leal, L. G. Migration of Rigid Spheres in a Two-Dimensional Unidirectional Shear Flow of a Second-Order Fluid. J. Fluid Mech. 1976, 76 (4), 783–799. 10.1017/S002211207600089X.

(51) D’Avino, G.; Romeo, G.; Villone, M. M.; Greco, F.; Netti, P. A.; Maffettone, P. L. Single Line Particle Focusing Induced by Viscoelasticity of the Suspending Liquid: Theory, Experiments and Simulations to Design a Micropipe Flow-Focuser. Lab. Chip 2012, 12 (9), 1638. 10.1039/c2lc21154h.

(52) Wu, M.; Chen, C.; Wang, Z.; Bachman, H.; Ouyang, Y.; Huang, P.-H.; Sadovsky, Y.; Huang, T. J. Separating Extracellular Vesicles and Lipoproteins *via* Acoustofluidics. Lab. Chip 2019, 19 (7), 1174–1182. 10.1039/C8LC01134F.

(53) Abreu, C. M.; Costa-Silva, B.; Reis, R. L.; Kundu, S. C.; Caballero, D. Microfluidic Platforms for Extracellular Vesicle Isolation, Analysis and Therapy in Cancer. Lab. Chip 2022, 22 (6), 1093–1125. 10.1039/D2LC00006G.

(54) Liu, C.; Guo, J.; Tian, F.; Yang, N.; Yan, F.; Ding, Y.; Wei, J.; Hu, G.; Nie, G.; Sun, J. Field-Free Isolation of Exosomes from Extracellular Vesicles by Microfluidic Viscoelastic Flows. ACS Nano 2017, 11 (7), 6968–6976. 10.1021/acsnano.7b02277.

(55) Johnson, D. W.; Goettert, J.; Singh, V.; Yemane, D. SUEX Dry Film Resist – A New Material for High Aspect Ratio Lithography.

(56) Izbassarov, D.; Rosti, M. E.; Ardekani, M. N.; Sarabian, M.; Hormozi, S.; Brandt, L.; Tammisola, O. Computational Modeling of Multiphase Viscoelastic and Elastoviscoplastic Flows. Int. J. Numer. Methods Fluids 2018, 88 (12), 521–543. 10.1002/fld.4678.

(57) Breugem, W.-P. A Second-Order Accurate Immersed Boundary Method for Fully Resolved Simulations of Particle-Laden Flows. J. Comput. Phys. 2012, 231 (13), 4469–4498. 10.1016/j.jcp.2012.02.026.

(58) Oldroyd, J. G.; Wilson, A. H. On the Formulation of Rheological Equations of State. Proc. R. Soc. Lond. Ser. Math. Phys. Sci. 1950, 200 (1063), 523–541. 10.1098/rspa.1950.0035.

(59) Hagey, D. W.; Ojansivu, M.; Bostancioglu, B. R.; Saher, O.; Bost, J. P.; Gustafsson, M. O.; Gramignoli, R.; Svahn, M.; Gupta, D.; Stevens, M. M.; Görgens, A.; El Andaloussi, S. The Cellular Response to Extracellular Vesicles Is Dependent on Their Cell Source and Dose. Sci. Adv. 2023, 9 (35), eadh1168. 10.1126/sciadv.adh1168.

(60) Corso, G.; Mäger, I.; Lee, Y.; Görgens, A.; Bultema, J.; Giebel, B.; Wood, M. J. A.; Nordin, J. Z.; Andaloussi, S. E. Reproducible and Scalable Purification of Extracellular Vesicles Using Combined Bind-Elute and Size Exclusion Chromatography. Sci. Rep. 2017, 7 (1), 11561. 10.1038/s41598-017-10646-x.

(61) Görgens, A.; Corso, G.; Hagey, D. W.; Jawad Wiklander, R.; Gustafsson, M. O.; Felldin, U.; Lee, Y.; Bostancioglu, R. B.; Sork, H.; Liang, X.; Zheng, W.; Mohammad, D. K.; Van De Wakker, S. I.; Vader, P.; Zickler, A. M.; Mamand, D. R.; Ma, L.; Holme, M. N.; Stevens, M. M.; Wiklander, O. P. B.; El Andaloussi, S. Identification of Storage Conditions Stabilizing Extracellular Vesicles Preparations. J. Extracell. Vesicles 2022, 11 (6), e12238. 10.1002/jev2.12238.

