## Supporting information for "Sheathless elasto-inertial focusing of sub-25 nm particles in straight microchannels"

Selim Tanriverdi^1^, Javier Cruz^1,2^, Shahriar Habibi^3^, Taras Sych^4^, Martim Costa^1^, Gustaf Mårtensson^1^, André Görgens^5,6,7^, Samir EL Andaloussi^5,6,7^, Luca Brandt^3,8^, Outi Tammisola^3^, Erdinc Sezgin^4^ & Aman Russom^1,9,*^

^1^ Division of Nanobiotechnology, Department of Protein Science, Science for Life Laboratory, KTH Royal Institute of Technology, Solna, 171 65, Sweden.

^2^ Division of Microsystems Technology, Department of Materials Science and Engineering, Uppsala University, Uppsala, 752 37, Sweden

^3^ FLOW and SeRC (Swedish e-Science Research Centre), Department of Engineering Mechanics, Royal Institute of Technology, Stockholm, SE 100 44, Sweden

^4^ Science for Life Laboratory, Department of Women’s and Children’s Health, Karolinska Institutet, Solna, Sweden

^5^ Division of Biomolecular and Cellular Medicine, Department of Laboratory Medicine, Karolinska Institutet, ANA Futura, Alfred-Nobels-Allé 8, 14152 Huddinge, Stockholm, Sweden

^6^ Department of Cellular Therapy and Allogeneic Stem Cell Transplantation (CAST), Karolinska University Hospital, 141 86 Stockholm, Sweden

^7^ Karolinska ATMP Center, Karolinska Institutet, ANA Futura, Alfred-Nobels-Allé 8, 14152 Huddinge, Stockholm, Sweden

^8^ Department of Environment, Land and Infrastructure Engineering, Politecnico di Torino, Turin, Italy

^9^ AIMES-Center for the Advancement of Integrated Medical and Engineering Sciences at Karolinska Institutet and KTH Royal Institute of Technology, Stockholm, Sweden

^*^

**Rheological Properties of PEO**

|  | **PEO Concentration (ppm)** | | | |
| --- | --- | --- | --- | --- |
|  | **500** | **1000** | **2000** | **4000** |
| **Density (kg/m^3^)** | 996 | 996 | 996 | 996 |
| **Zero-shear viscosity µ_0_ (mPa.s)** | 1.26 | 1.59 | 2.39 | 8.23 |
| **Effective relaxation time 𝝀_e_ (ms)** | 4.31 | 6.76 | 10.61 | 16.65 |
| **Overlap concentration c^*^ (ppm)** | 858 | 858 | 858 | 858 |

Table S1. The rheological properties of the viscoelastic fluid (PEO M_W_ = 2×10^6^g/mol)

**Dimensionless Numbers**

| **Q (µL/min)** | **PEO Concentration (ppm)** | | | | | | | | | | | |
| --- | --- | --- | --- | --- | --- | --- | --- | --- | --- | --- | --- | --- |
|  | **500** | | | **1000** | | | **2000** | | | **4000** | | |
|  | **Re** | **Wi** | **El** | **Re** | **Wi** | **El** | **Re** | **Wi** | **El** | **Re** | **Wi** | **El** |
| 0.5 | 0.20 | 48 | 236 | 0.16 | 76 | 470 | 0.11 | 118 | 1102 | 0.03 | 185 | 5955 |
| 1 | 0.41 | 96 | 236 | 0.32 | 151 | 470 | 0.21 | 236 | 1102 | 0.06 | 370 | 5955 |
| 1.5 | 0.61 | 143 | 236 | 0.48 | 227 | 470 | 0.32 | 353 | 1102 | 0.09 | 555 | 5955 |
| 2 | 0.81 | 191 | 236 | 0.64 | 302 | 470 | 0.43 | 471 | 1102 | 0.12 | 740 | 5955 |
| 2.5 | 1.01 | 239 | 236 | 0.80 | 378 | 470 | 0.53 | 589 | 1102 | 0.16 | 925 | 5955 |
| 3 | 1.22 | 287 | 236 | 0.96 | 453 | 470 | 0.64 | 707 | 1102 | 0.19 | 1110 | 5955 |

Table S2. Dimensionless numbers in microchannel with height=60 µm and width=5 µm

| **Q (µL/min)** | **PEO Concentration (ppm)** | | | | | | | | | | | |
| --- | --- | --- | --- | --- | --- | --- | --- | --- | --- | --- | --- | --- |
|  | **500** | | | **1000** | | | **2000** | | | **4000** | | |
|  | **Re** | **Wi** | **El** | **Re** | **Wi** | **El** | **Re** | **Wi** | **El** | **Re** | **Wi** | **El** |
| 0.5 | 0.19 | 12 | 63 | 0.15 | 19 | 127 | 0.1 | 29 | 297 | 0.03 | 46 | 1603 |
| 1 | 0.38 | 24 | 63 | 0.3 | 38 | 127 | 0.2 | 59 | 297 | 0.06 | 93 | 1603 |
| 1.5 | 0.56 | 36 | 63 | 0.45 | 57 | 127 | 0.3 | 88 | 297 | 0.09 | 139 | 1603 |
| 2 | 0.75 | 48 | 63 | 0.6 | 76 | 127 | 0.4 | 118 | 297 | 0.12 | 185 | 1603 |
| 2.5 | 0.94 | 60 | 63 | 0.75 | 94 | 127 | 0.5 | 147 | 297 | 0.14 | 231 | 1603 |
| 3 | 1.13 | 72 | 63 | 0.89 | 113 | 127 | 0.6 | 177 | 297 | 0.17 | 278 | 1603 |

Table S3. Dimensionless numbers in microchannel with height=60 µm and width=10 µm

**Effect of Channel Width**

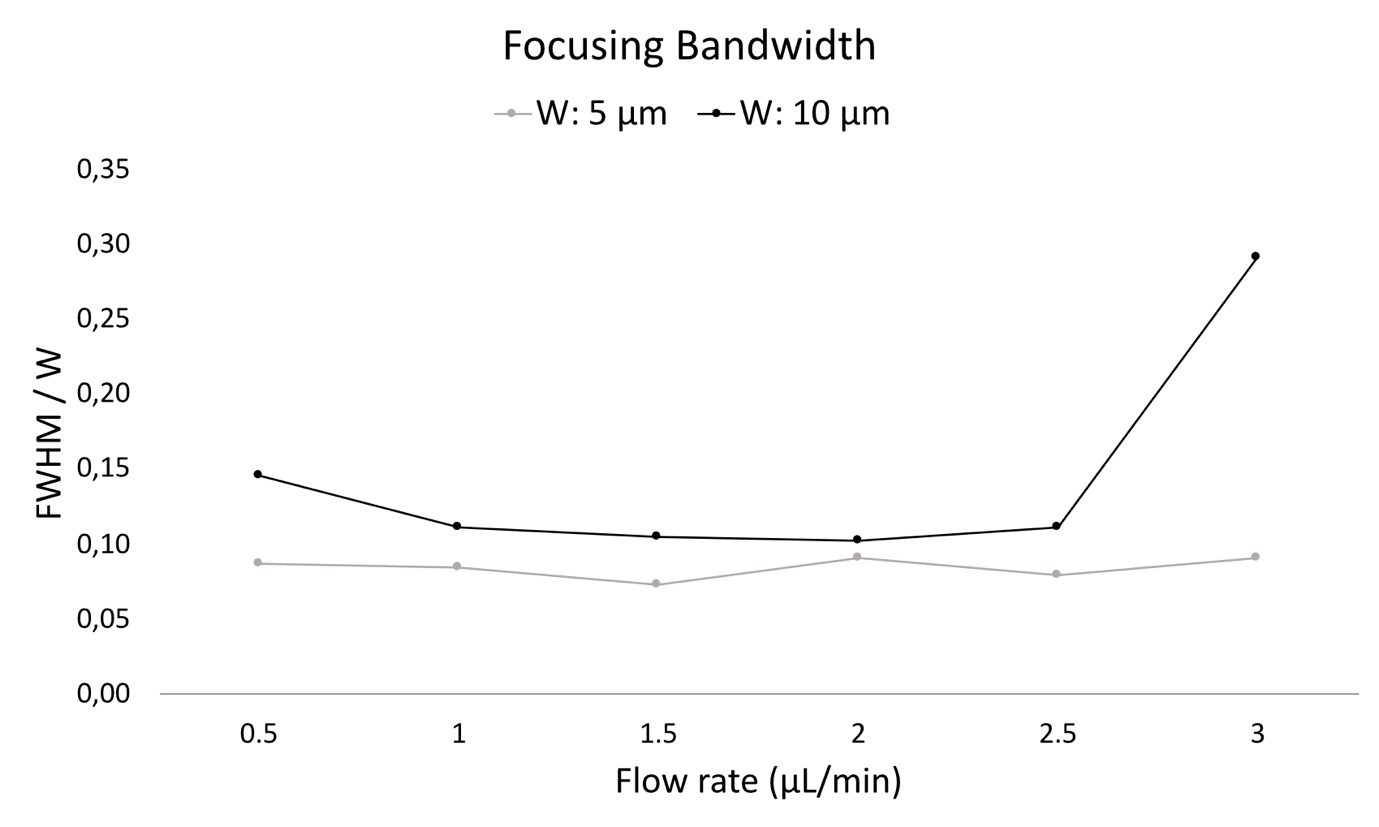

Figure S1. Focusing bandwidth comparison of w=5 µm and 10 µm for 50 nm particles at 1000 ppm PEO concentration

**Numerical Analysis of Aspect Ratio**

Our objective is to model the migration of particles in viscoelastic (VE) fluids within a straight microchannel characterized by a relatively high aspect ratio. We examine the effect of the aspect ratio of the microchannel and particle sizes on the migration of particles. Specifically, particles with diameters of 1.6 𝜇𝑚, 1 𝜇𝑚, and 750 𝑛𝑚 are studied under identical conditions of constant flow rate and fluid rheological properties. For simulations with an aspect ratio 𝐴𝑅 = 12, we employ a channel with dimensions of 60 𝜇𝑚 in height and 5 𝜇𝑚 in width. In contrast, for simulations with an 𝐴𝑅 = 6, the channel dimensions are adjusted to 30 𝜇𝑚 in height while maintaining a width of 5 𝜇𝑚. We expect to observe different particle migration patterns among these different aspect ratios and particle sizes.

The migration behaviour of the particle with a diameter of 1.6 𝜇𝑚 for two different aspect ratios of 6 and 12 is illustrated in Figure S2(b). We observe that in both microchannels, despite their differing aspect ratios, particles ultimately reach the centerline. This behaviour indicates the predominance of elastic forces over inertial forces in the fluid. However, the particle has a faster migration toward the centerline in the microchannel with 𝐴𝑅 = 12 compared to the channel with 𝐴𝑅 = 6. The observed behaviour can be attributed to the previously mentioned relationship between aspect ratio and normal stress difference. Specifically, a higher aspect ratio leads to an increased normal stress difference in the fluid. This enhancement in stress difference translates to a stronger force acting on the particle’s surface. Consequently, particles in channels with higher aspect ratios experience a more pronounced push towards the equilibrium position at the channel centerline.

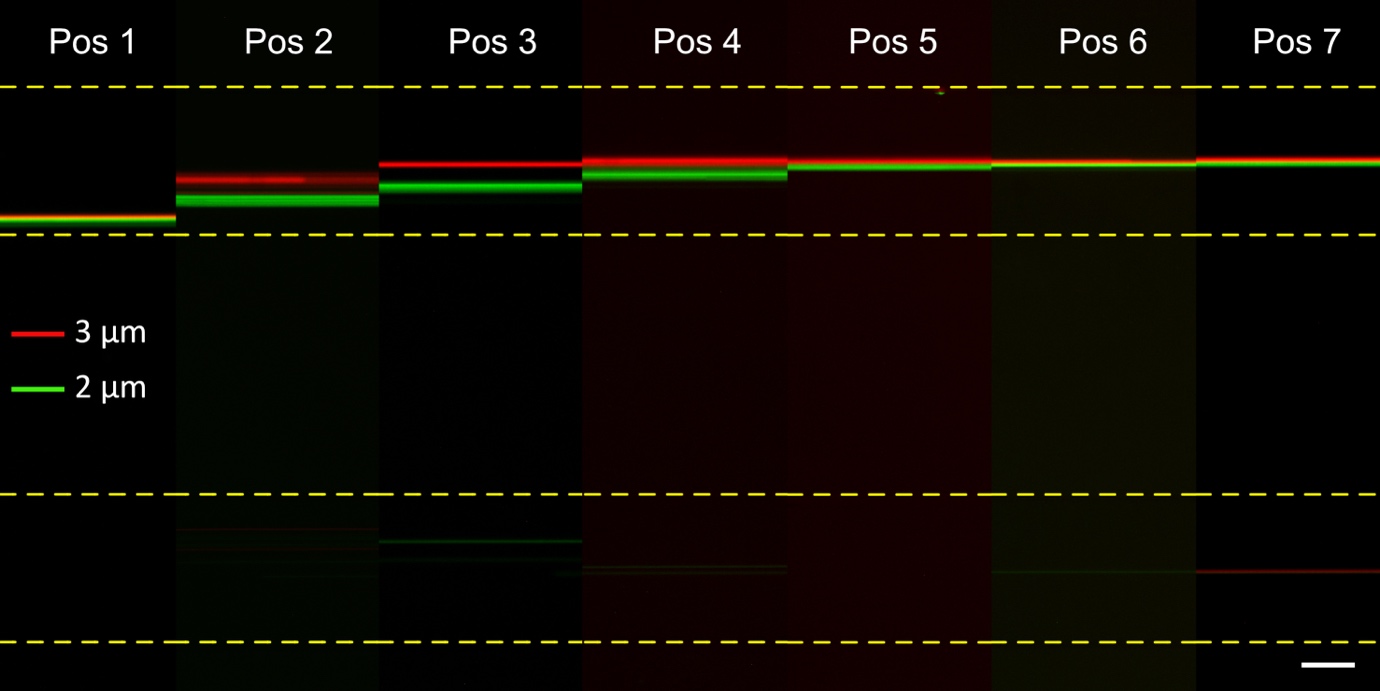

Figure S3. Migration of 2 µm and 3 µm particles for size-based particle separation. Scale bar: 50 µm.

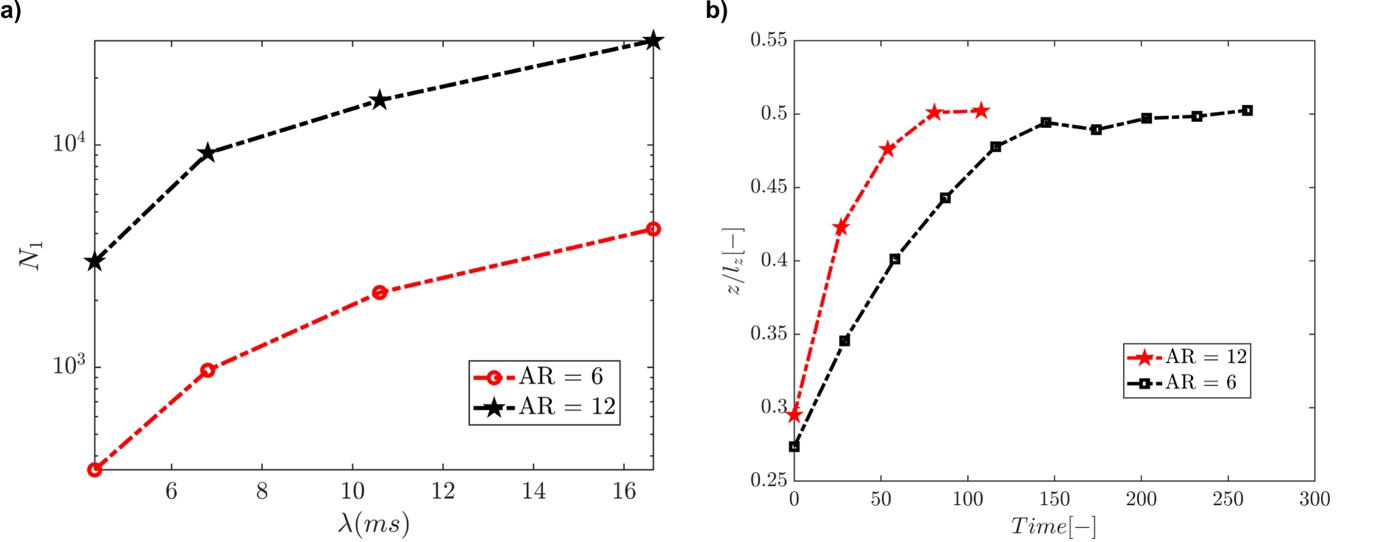

Figure S2. The numerical comparison of AR=6 and 12 – a) The variation of the first normal stress difference across the cross–section of the channel regarding different relaxation times b) Particle spanwise position over time

**Size Distribution of Biological Nanoparticles**

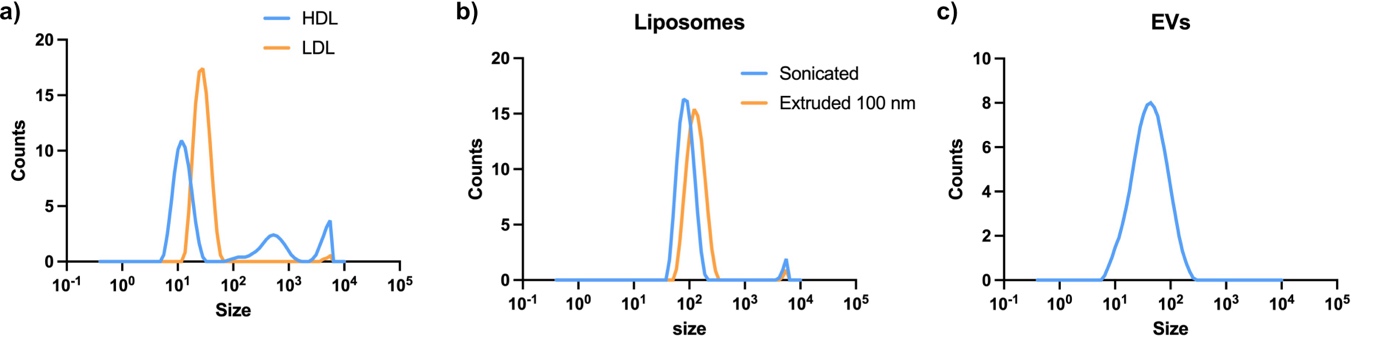

Figure S4. Size distribution of a) HDL–LDL b) Liposomes and c) EVs
